# Environmental pH impacts division assembly and cell size in *Escherichia coli*

**DOI:** 10.1101/747808

**Authors:** Elizabeth A. Mueller, Corey S. Westfall, Petra Anne Levin

## Abstract

Cell size is a complex trait, derived from both genetic and environmental factors. Environmental determinants of bacterial cell size identified to date primarily target assembly of cytosolic components of the cell division machinery. Whether certain environmental cues also impact cell size through changes in the assembly or activity of extracytoplasmic division proteins remains an open question. Here, we identify extracellular pH as a modulator of cell division and a key determinant of cell size across evolutionarily distant bacterial species. In the Gram-negative model organism *Escherichia coli*, our data indicate environmental pH impacts the length at which cells divide by altering the ability of the terminal cell division protein FtsN to localize to the cytokinetic machinery and activate division. Acidic environments lead to enrichment of FtsN at the septum and activation of division at a reduced cell length, while alkaline pH inhibits FtsN localization and suppress division activation. Altogether, our work reveals a previously unappreciated role for pH in bacterial cell size control.

## INTRODUCTION

A fundamental property of cells, size is tightly linked to physiological state. With few exceptions, two processes dictate cell size: cell growth and cell cycle progression. During steady state or “balanced” growth, bacteria add on average the same volume between cell birth and division regardless of their size at birth. This phenomenon, referred to as the ‘adder’ model for bacterial cell size homeostasis, results in convergence to an average cell size [1,2]. Simulations and experimental data suggest that the adder is an emergent property of two processes: 1) growth rate dependent synthesis of rate-limiting components of the cell division machinery, and 2) accumulation of these proteins to threshold numbers necessary to support cytokinesis. Consistent with this model, perturbing the normal accumulation of one such protein, the tubulin homolog FtsZ, disrupts the volume added between divisions. Although sufficient to alter size at the onset of DNA replication, disruptions in levels of the DNA replication initiation protein DnaA fail to impact homeostatic cell size, indicating that cell division—and not cell cycle progression generally—ultimately controls the volume of new material cells add during steady state growth [3].

Although there is little variation in size during steady state growth under a single, constant condition, changes in the environment can drastically affect average cell size. Nutrient availability, in particular, has a dramatic and positive impact on the size of evolutionarily distant bacteria, including *Escherichia coli*, *Salmonella enterica*, and *Bacillus subtilis*, and single-celled yeasts [4–7]. Bacterial cells amass up to three times the volume when cultured in nutrient rich conditions as compared to nutrient poor conditions. Cell length and cell width both scale with nutrient availability in *E. coli* [8], while width remains nearly constant for *B. subtilis* [6]. The molecular basis of the positive relationship between nutrients and cell size is multifactorial, involving changes in biosynthetic processes that underlie cell growth [9,10] and cell cycle progression [6,7,11]. Notably, nutrient-dependent changes in cell cycle progression identified to date all impinge on FtsZ assembly. In *B. subtilis* and *E. coli*, accumulation of the metabolite uridine disphosphate (UDP)-glucose during growth in carbon rich media activates two unrelated glucosyltransferases, UgtP and OpgH, which antagonize FtsZ ring assembly. Although mechanistically distinct, both antagonists functionally increase the threshold quantity of FtsZ that must accumulate prior to cytokinesis. In *E. coli*, the threshold is almost certainly further heightened through nutrient-dependent changes in width, which require the construction of a larger cytokinetic ring.

While regulatory mechanisms coordinating division, nutrient availability, and size are well documented, comparatively little is known about how the cell division machinery responds to other environmental cues. The division machinery in *E. coli* consists of over 20 proteins, collectively referred to the divisome. These proteins assemble in a hierarchical fashion, beginning with mid-cell polymerization of FtsZ in the cytosol and ending with recruitment of extracytoplasmic septal cell wall synthesis enzymes and their regulators [12,13]. Exterior to the plasma membrane, these so-called ‘late’ division proteins are directly exposed to dynamic and potentially extreme environmental conditions—including changes in pH, osmolarity, and ionic strength—that may impact their ability to activate and complete cross wall synthesis [14,15]. Differential activation of the divisome has the potential to impact cell size homeostasis: hypermorphic mutants of *E. coli* and *Caulobacter crescentus* that affect the initiation or rate of septal cell wall synthesis are consistently short independent of changes to growth rate [16–20]. However, whether these represent native integration points for environmental modulation of cell size homeostasis remains unclear.

Here, we identify environmental pH as a conserved, growth rate independent determinant of cell size in evolutionarily distant bacterial species. Distinct from nutrient-dependent changes in size, pH does not alter FtsZ assembly. Instead, pH-dependent changes in *E. coli* cell length appear to stem from differential localization of the terminal division protein and cell wall synthesis activator, FtsN. Collectively, our data support a model in which pH-dependent changes in accumulation of FtsN at the cytokinetic ring impact the volume at which cells initiate division, thereby altering average cell size.

## RESULTS

### pH influences cell size in diverse bacteria

To investigate the impact of pH on cell size, we cultured *E. coli* strain MG1655 at steady state in nutrient rich media (LB + 0.2% glucose) under a physiologically relevant range of pH conditions (pH 4.5-8.5) [21,22]. We sampled cultures and fixed the cells during early exponential phase (OD_600_ ∼ 0.1-0.2) for cell size analysis; at this time point, the pH of the culture had not significantly deviated from the starting pH (SI Appendix, Figure S1). Strikingly, cells cultured at pH 4.5 were ∼75% of the area of their counterparts grown at pH 7.0. In contrast, cells cultured at pH 8.5 were ∼120% of the area of cells grown at pH 7.0 (Figure 1A, B; SI Appendix, Figure S2 and Table S3). Apart from the most extreme acidic conditions, nearly all of the pH-dependent changes in size were restricted to changes in cell length and were independent of changes in growth rate, media composition, or buffering capacity (Figure 1C, D; SI Appendix, Figure S1, Table S3). To independently validate pH alters cell size homeostasis in live cells, we used time lapse imaging to measure the cell length added between divisions, a property of the adder model of cell size homeostasis [1,2]. Consistent with our findings in fixed cells, cells cultured in acidic media added a shorter length from birth to division than cells grown at neutral and alkaline conditions (Figure 1E).

**Figure 1.**
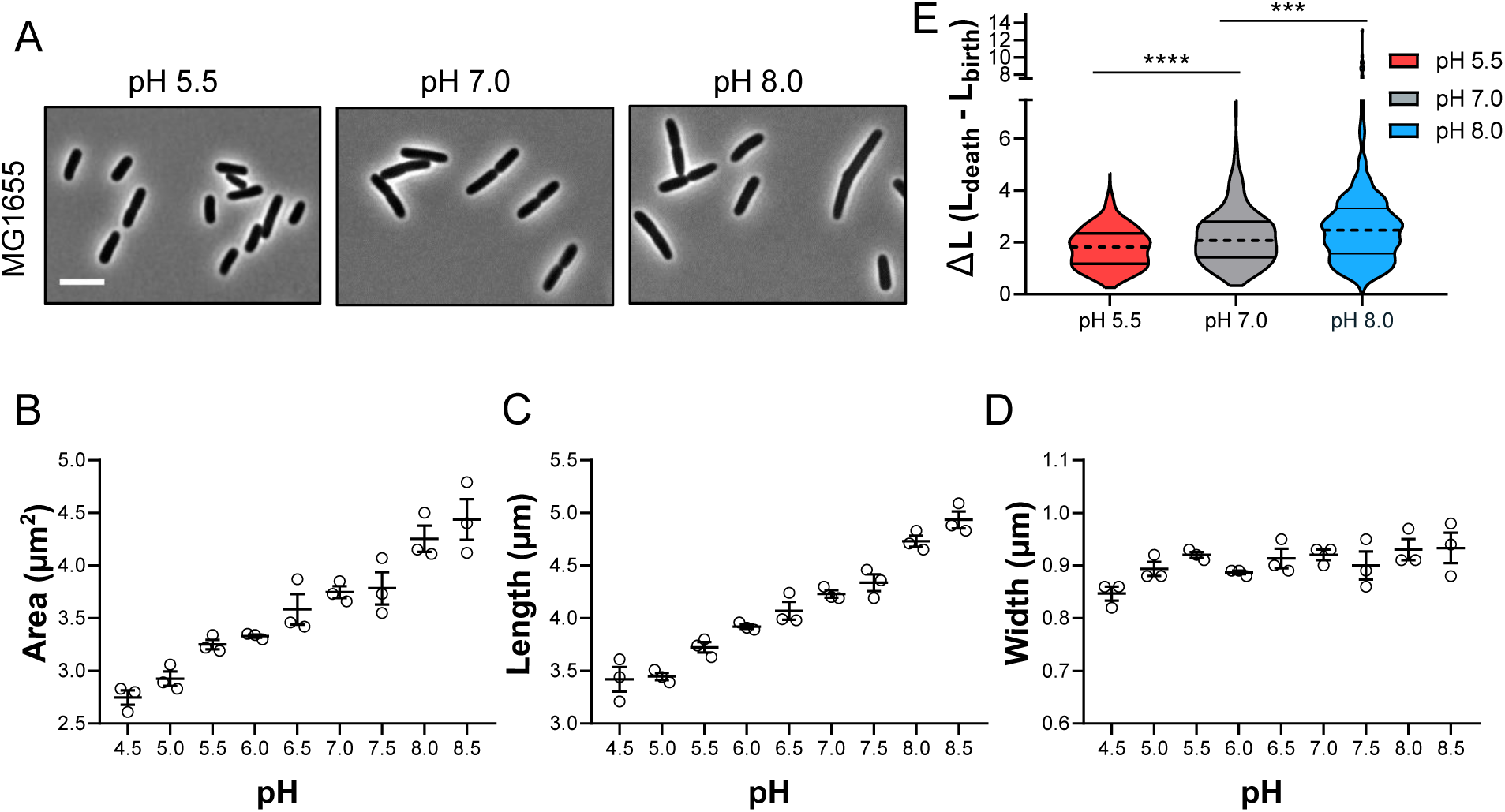
Environmental pH influences *E. coli* cell size. A) Representative micrographs of MG1655 grown to steady state in LB media + 0.2% glucose at pH 5.5, 7.0, and 8.0. Scale bar denotes 5 µm. B-D) Mean cell area (B), cell length (C), and cell width (D) for MG1655 grown in LB media + 0.2% glucose from pH 4.5-8.5. Individual points denote mean population measurement for each biological replicate. Error bars represent standard error of the mean. Significance shown in SI Appendix, Table S3 E) Change in length from beginning to end of cell cycle for individual cells grown in LB media + 0.2% glucose at pH 5.5 (n = 450), pH 7.0 (n = 489), or pH 8.0 (n = 461) from two independent experiments. Dotted line represents median length added, and straight lines indicate quartiles. Significance was determined by Kruskal-Wallis text, corrected for multiple comparisons with Dunn’s test.

pH-dependent changes in size were not unique to MG1655 or even to *E. coli*. We observed similar effects of pH on cell area in *E. coli* strain W3110 and in the evolutionary distant Gram-positive coccus *Staphylococcus aureus* (SI Appendix, Figure S3). Likewise, during this work two separate studies noted the size of *Streptococcus pneumoniae* and *C. crescentus* also increases during growth in alkaline media [23]. Altogether, these findings establish environmental pH as mediator of size across evolutionary distant bacterial species.

### Acidic pH stabilizes late division proteins and bypasses the essentiality of FtsK

Our observation that pH-dependent changes in *E. coli* cell size were restricted to changes in cell length and were independent of changes in mass doubling time (Figure 1C; SI Appendix, Table S3) indicated divisome assembly and/or activity may be pH sensitive. In *E. coli* assembly of the ‘core’ cell division machinery is a sequential process [12]. First, the so-called “early” division proteins—including the cytosolic tubulin homolog FtsZ, membrane anchor ZipA, membrane-associated actin homolog FtsA—form a dynamic, discontinuous ring-like structure at a mid-cell [24–26]. Subsequently, a series of “late” division proteins containing transmembrane periplasmic domains becomes enriched at the septal ring; these include the DNA translocase FtsK [27,28], the regulatory FtsQLB complex [18,19,29], and septal PG synthesis transpeptidase and glycosyltransferase FtsI and FtsW [30]. In the final phase of division in proteobacteria, FtsN accumulates at mid-cell and is believed to “trigger” septal PG synthesis and constriction through activation of early and late divisome components [18,19,31–33]. In addition to the essential division proteins described above, there are nearly a dozen non-essential or conditionally essential factors involved in divisome stabilization (e.g., ZapBCD and FtsP) [34–37], cell wall synthesis (e.g., PBP1a and PBP1b) [38,39], cell wall hydrolysis (e.g., AmiA and AmiC) [40], and regulation of cell wall remodeling (e.g., FtsEX) [41,42].

The sheer number of proteins involved in division imply many possible integration points through which pH may modulate division to tune cell size. Based on our previous finding that pH impacts the activity of periplasmic cell wall synthases [43], we speculated that division proteins with periplasmic domains would be the most likely regulatory targets of pH. Located between *E. coli’s* plasma membrane and its semi-permeable outer membrane, the periplasm is particularly sensitive to changes in the abundance of ions and other small molecules and assumes the environmental pH while the cytosol remains relatively buffered at steady state [14][15].

To identify the specific stage(s) of cell division influenced by pH, we took advantage of a set of heat sensitive alleles of essential cell division genes. These conditional mutants played a historically important role in parsing the key functions of the essential components of divisome and associated modulatory proteins [44,45]. Suppression of the heat sensitive phenotype of these conditional mutants under growth restrictive conditions (LB-no salt, 37 or 42°C) suggests a positive influence on the division machinery while enhancement of heat sensitivity under growth permissive conditions (LB, 30°C) indicates a negative influence on the division machinery.

We assessed the impact of pH on the growth of a subset of heat-sensitive mutants, including alleles of both early division genes [*ftsZ84* (G105S) and *ftsA27* (S195P] and late division genes [*ftsK44* (G80A), *ftsQ1* (E125K), and *ftsI23* (Y380D)]. Although growth of the early division mutants was insensitive to pH, low pH (5.5) suppressed the heat sensitivity of the late division mutants while high pH (8.0) enhanced it (Figure 2A). Importantly, this effect was not allele-specific. Additional heat-sensitive variants of FtsZ (*ftsZ25*), FtsA (*ftsA12*), and FtsI (*ftsI2158*) behaved similarly to their previously tested cognates with one exception: *ftsA12* heat sensitivity was modestly enhanced at pH 8.0 (SI Appendix, Figure S5). When we expanded the tested pH range from pH 4.5-9.0, the heat sensitivity of the strains encoding *ftsK44*, *ftsQ1*, and *ftsI23* was consistently suppressed between pH 5.0 and pH 6.5 and enhanced between pH 7.5 and pH 9.0 (SI Appendix, Figure S5). Notably, these pH ranges correspond to conditions in which the wild type cells have decreased and increased average cell lengths, respectively (Figure 1C). We did not observe changes in heat sensitivity in strains encoding *ftsZ84* or *ftsA27* in the expanded pH range (Figure 2B; SI Appendix, Figure S5). We also ruled out the contribution of the accessory periplasmic divisome proteins FtsP, PBP1a, and PBP1b, which had been previously shown to be stress or pH responsive [37,43]; cells defective in each of the aforementioned proteins still exhibited pH-dependent changes in cell size (SI Appendix, Figure S4).

**Figure 2.**
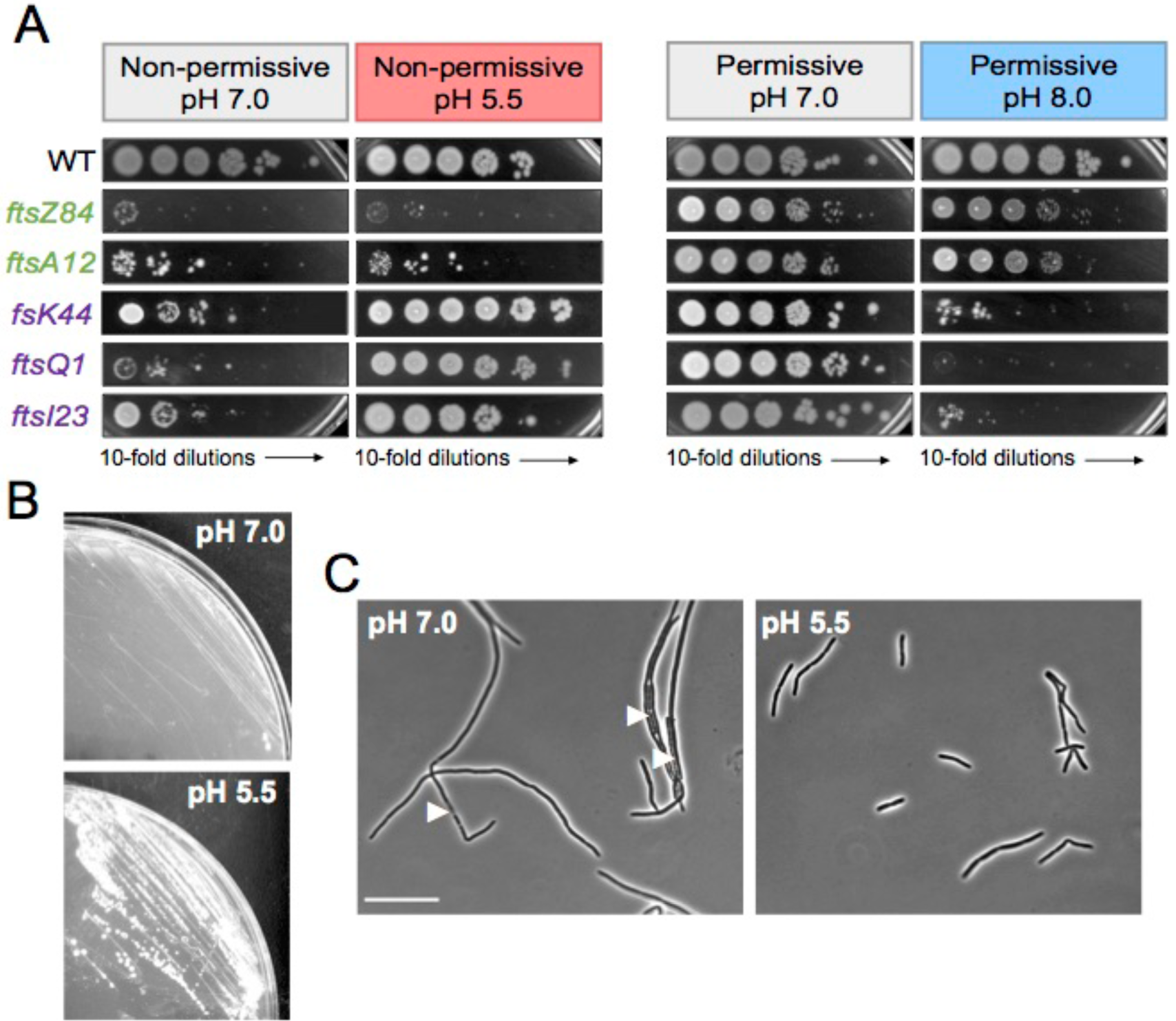
Acidic pH stabilizes late division proteins and bypasses the essentiality of FtsK. A) Representative plating efficiency for strains producing heat sensitive variants of early and late division proteins when grown under non-permissive conditions (left) or permissive conditions (right) as a function of agar plate pH. Image is representative of three biological replicates. B) Comparison of growth of MG1655 Δ*ftsK*::*kan* strain cultured on LB agarose plate at neutral (top) or acidic pH (bottom) at 30 °C. C) Comparison of cell morphology of MG1655 Δ*ftsK*::*kan* strain grown for 2 hours in LB liquid media at neutral (left) or acidic (pH). Arrows indicate lysed cells. Scale bar denotes 20 µm.

The complete suppression of heat sensitivity in *ftsK44* and *ftsI23* mutants at very high temperature (42 °C) suggested these genes may be dispensable for divisome activity in acidic media. To test this, we attempted to transduce deletion alleles of each gene into wild type cells under acidic (pH 5.5) and neutral conditions at 30 °C. Although we were unable to delete the native *ftsI* even in the presence of a catalytically inactive variant of FtsI (S307A) produced from a plasmid [46], we were able generate stable transductants with the *ftsK*::*kan* allele when the cells were grown in acidic media (Figure 2B). FtsK null mutants were slightly elongated when cultured in acidic media but rapidly filamented and lysed when transferred to neutral pH (Figure 2C). In total, these findings are consistent with acidic pH stabilizing one or more late division proteins, and this is sufficient to bypass the essential activity of FtsK.

### Septal recruitment of the terminal division protein FtsN is pH sensitive

To directly visualize the effect of pH on the assembly of the division machinery, we imaged mid-cell recruitment of a subset of GFP-tagged division proteins. We selected fusion proteins that spanned the divisome assembly hierarchy, including “early” cytoplasmic proteins FtsZ and FtsA, “late” periplasmic proteins FtsL and FtsI, and the terminal periplasmic division protein FtsN (Figure 3A). All constructs were integrated at the lambda locus in otherwise wild type MG1655 cells with the gene of interest expressed from an IPTG-inducible promoter. Production of these fusion proteins leads to a characteristic mid-cell ring of fluorescence for a fraction of the cell cycle proportional to their lifetime at the septum [47]. Because populations of *E. coli* cell are unsynchronized and thus all cell cycle stages are represented at a single time point, comparison of septal ring frequency across conditions can be used as a proxy for changes in assembly dynamics and/or enrichment of division proteins at mid-cell [47]. IPTG levels were titrated such that fusion protein production did not disrupt pH-dependent changes in size (SI Appendix, Figure S6).

**Figure 3.**
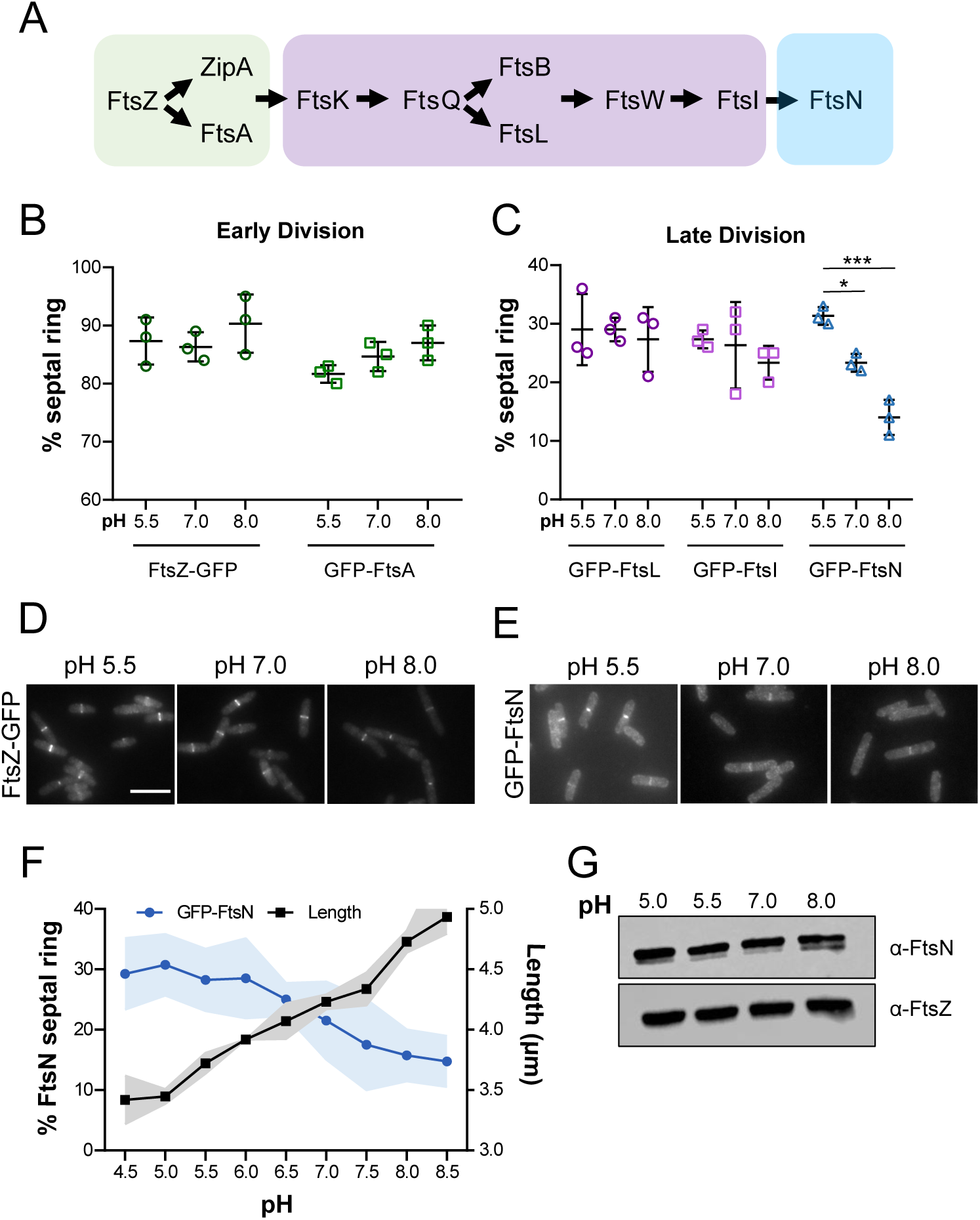
Septal recruitment of the terminal division protein FtsN is pH sensitive. A) Schematic depicting recruitment hierarchy of the early division proteins (green), late division proteins (purple), and final division protein FtsN (blue) to the mid-cell. B-C) Mean percentage of cells that score positive for a septal ring of the indicated early (left) and late (right) GFP-tagged division proteins in LB media at pH 5.5, 7.0, and 8.0. Individual points depict values of individual biological replicates. Error bars represent standard deviation. Significance was determined using a two-way ANOVA, corrected for multiple comparisons with Sidak’s test. D-E) Representative micrographs of cells with FtsZ-GFP rings (left) and GFP-FtsN rings (right) at pH 5.5, 7.0, and 8.0. Scale bar denotes 5 µm. F) Comparison of mean FtsN septal ring frequency and mean cell length from pH 4.5-8.5. Cell length data is from Figure 1C. Shaded region denotes the error of the measurement (SD for ring frequency and SEM for cell length). G) Representative immunoblot for FtsN and FtsZ levels in MG1655 in LB media at pH 5.0, 5.5, 7.0, and 8.0. Quantification shown in SI Appendix, Figure S8.

When we compared the septal ring frequency of the aforementioned fusion proteins at pH 5.5, 7.0, and 8.0, only cells producing GFP-FtsN exhibited pH-dependent changes in mid-cell localization of the protein (Figure 3; SI Appendix, Table S4). GFP-FtsN was significantly enriched at the mid-cell under acidic conditions (∼30% septal localization frequency) and reduced under alkaline conditions (∼15% septal localization frequency). This trend held across an expanded pH range (4.5-8.5) and was inversely proportional to changes in cell length (Figure 3D). To validate the septal ring frequency analysis, we quantified mid-cell GFP-FtsN intensity across pH conditions using the software Coli Inspector, which confirmed specific enrichment of FtsN at mid-cell at low pH (SI Appendix, Figure S7) [47].

Two models explain pH-dependent changes in mid-cell localization of FtsN: 1) increased expression, production, or stability of FtsN in acidic conditions and/or 2) changes in FtsN’s affinity for the cytokinetic machinery. To address the former possibility, we compared bulk levels of FtsN from cells grown in varying pH environments. Neither the levels of the native or GFP-tagged variant of FtsN varied as a function of pH by immunoblot (Figure 3E; SI Appendix, Figure S6 and S8), indicating the observed changes in ring frequency are likely due to an increase in affinity for the septal ring under acidic conditions.

### Overexpression of FtsN decreases cell length

If FtsN localization to the otherwise mature divisome is sufficient to trigger constriction, enhancing interaction between FtsN and other components of the division machinery by overexpression should lead to reductions in cell length. To test this model, we overexpressed *gfp-ftsN* from a plasmid in the wild type background at neutral pH. Consistent with FtsN serving as a division “trigger”, we observed an induction-dependent decrease in cell length of up to ∼15%, which correlated with an increase in septal localization frequency (Figure 4; SI Appendix, Table S4). Cell width did not decrease upon FtsN overexpression and in fact, modestly increased. These results are counter to previous work reporting toxicity and a modest increase in cell length with *ftsN* overexpression [12,45]. To confirm our findings, we repeated this experiment using a different expression construct and again observed a similar reduction in cell length (SI Appendix, Figure S10). Importantly, overexpression of other late division proteins has not been associated with reductions in cell size. Simultaneous overexpression of the *ftsQ*, *ftsL*, and *ftsB* causes cell filamentation [48], and in our hands overexpression of *ftsI* also modestly increases cell length (SI Appendix, Figure S10).

**Figure 4.**
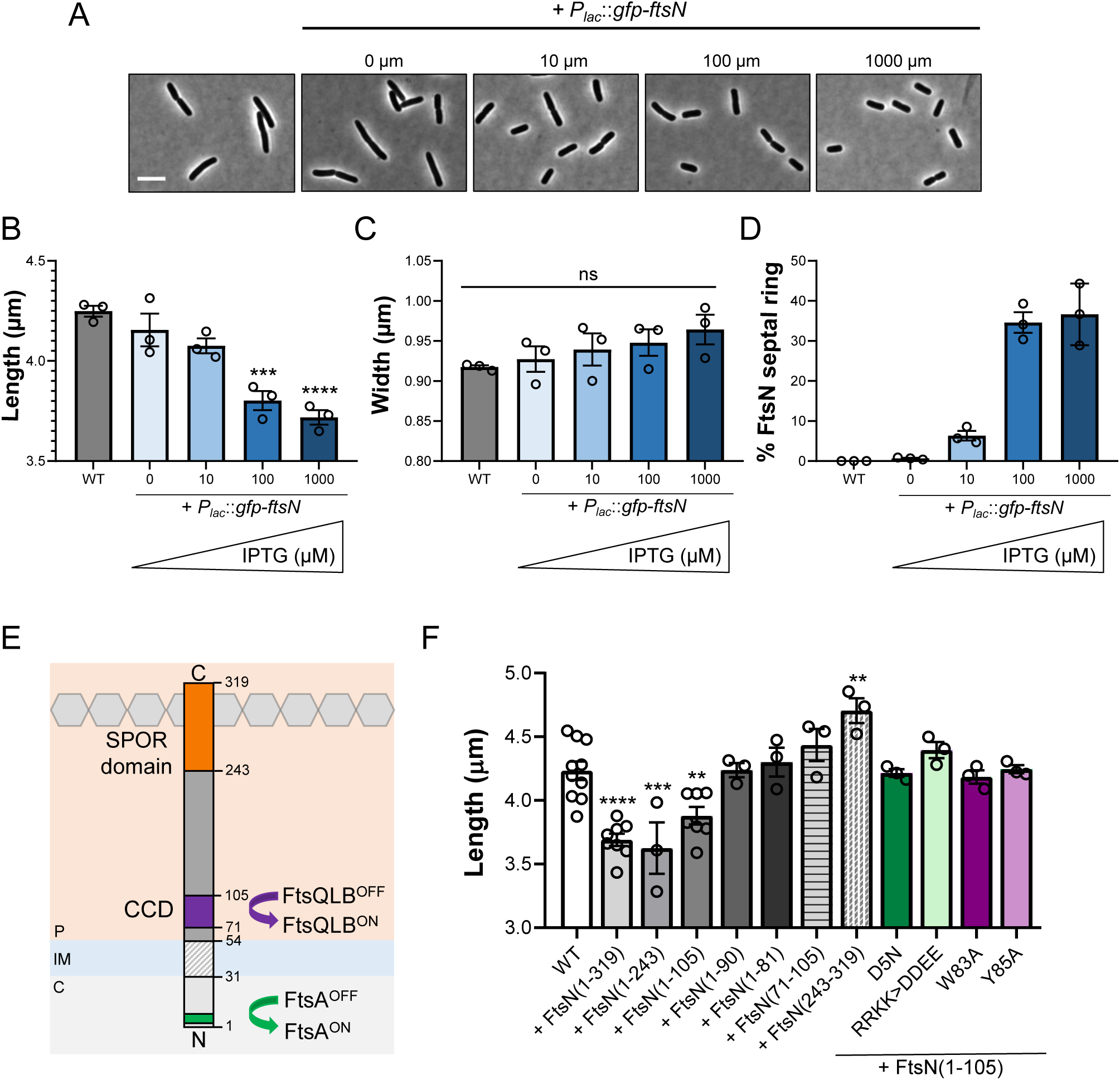
Overexpression of FtsN reduces cell length. A) Representative micrographs of MG1655 at varying levels of *gfp-ftsN* overexpression in LB media. Scale bar denotes 5 µm. B-D) Mean cell length (B), cell width (C), and FtsN septal ring frequency (D) of cells overexpressing *gfp-ftsN* grown in LB media. Individual points depict measurement from each biological replicate. Error bars represent standard error of the mean (B, C) or standard deviation (D). Significance was determined by a one-way ANOVA, normalized for multiple comparisons with Dunnett’s test. E) Schematic depicting the major features of the FtsN gene product, including an FtsA interaction interface (green), essential constriction control domain (CCD, purple), and peptidoglycan-binding SPOR domain (orange). F) Mean cell length of MG1655 overexpressing *gfp-ftsN* truncations and point mutants during growth in LB media. All strains harboring a construct were induced with 1 mM IPTG. Individual points depict mean population measurement from each biological replicate. Error bars represent standard error of the mean. Significance was determined by a one-way ANOVA, normalized for multiple comparisons with Dunnett’s test.

We next sought to determine the regions of FtsN that are necessary and sufficient for overexpression-dependent reductions in cell size. A bitopic inner membrane protein with no known enzymatic activity, FtsN is believed to activate division through two mechanisms: direct interaction with the early division protein FtsA through a short, N-terminal cytoplasmic region and genetic interaction with the FtsQLB complex through an essential alpha helical region in the periplasm referred to as the constriction control domain (CCD) [18,32,49]. Additionally, FtsN possesses a periplasmic C-terminal SPOR domain, which binds denuded glycans in the peptidoglycan cell wall and enhances FtsN septal localization after the onset of constriction initiation [32,50] (Figure 4E).

To identify the regions of FtsN that are sufficient to reduce length, we overexpressed a series of *ftsN* truncation mutants in wild type cells at neutral pH and measured their size [32]. Our data demonstrate the N-terminal 105 amino acids of FtsN, which include both the FtsA interaction interface and the CCD, are sufficient for overexpression-dependent reductions in cell length (Figure 4F). Overexpression of *ftsN* encoding only part of the CCD (1-81, 1-90), the CCD alone (71-105), or the SPOR domain alone (243-319) did not significantly reduce length despite being produced at similar levels (SI Appendix, Figure S11). Somewhat surprisingly, the SPOR domain was not required for reductions in length, suggesting a direct interaction with the PG is not necessary for FtsN’s role in size control [50]. Overexpression of *ftsN(1-105)* mutants impaired for FtsA interaction (D5A, RRKK>DDEE) or CCD activity (Y83A, W85A) failed to reduce length despite being stably produced (Figure 4F; SI Appendix, Figure S11). Collectively, this functional analysis establishes the FtsA interaction interface and the CCD as requirements for FtsN-mediated reductions in cell length.

### Acidic pH activates division through FtsN

Hypermorphic alleles of *ftsA*, *ftsL*, and *ftsB* have been identified that mimic the stimulatory effect of FtsN on the divisome and consequently bypass FtsN’s essential function. Cells expressing these hypermorphs are constitutively short, independent of changes in growth rate [16,18,19,42]. We reasoned that if environmental pH modulates division through activation of FtsA and FtsQLB, the size of cells expressing hypermorphic alleles would be insensitive to pH (Figure 5A). To test this model, we compared size and FtsN localization in two of most well-studied division hypermorphs, *ftsA\** (R286W) and *ftsL\** (E88K), at pH 5.5, 7.0, and 8.0. As expected, the size of *ftsA\** and *ftsL\** mutants was invariant across pH conditions (Figure 5B). At the same time, the frequency of septal GFP-FtsN was higher in the hypermorphic strains (41% and 38%, respectively, compared to 21% of wild type cells) (Figure 5C; SI Appendix, Figure S12).

**Figure 5.**
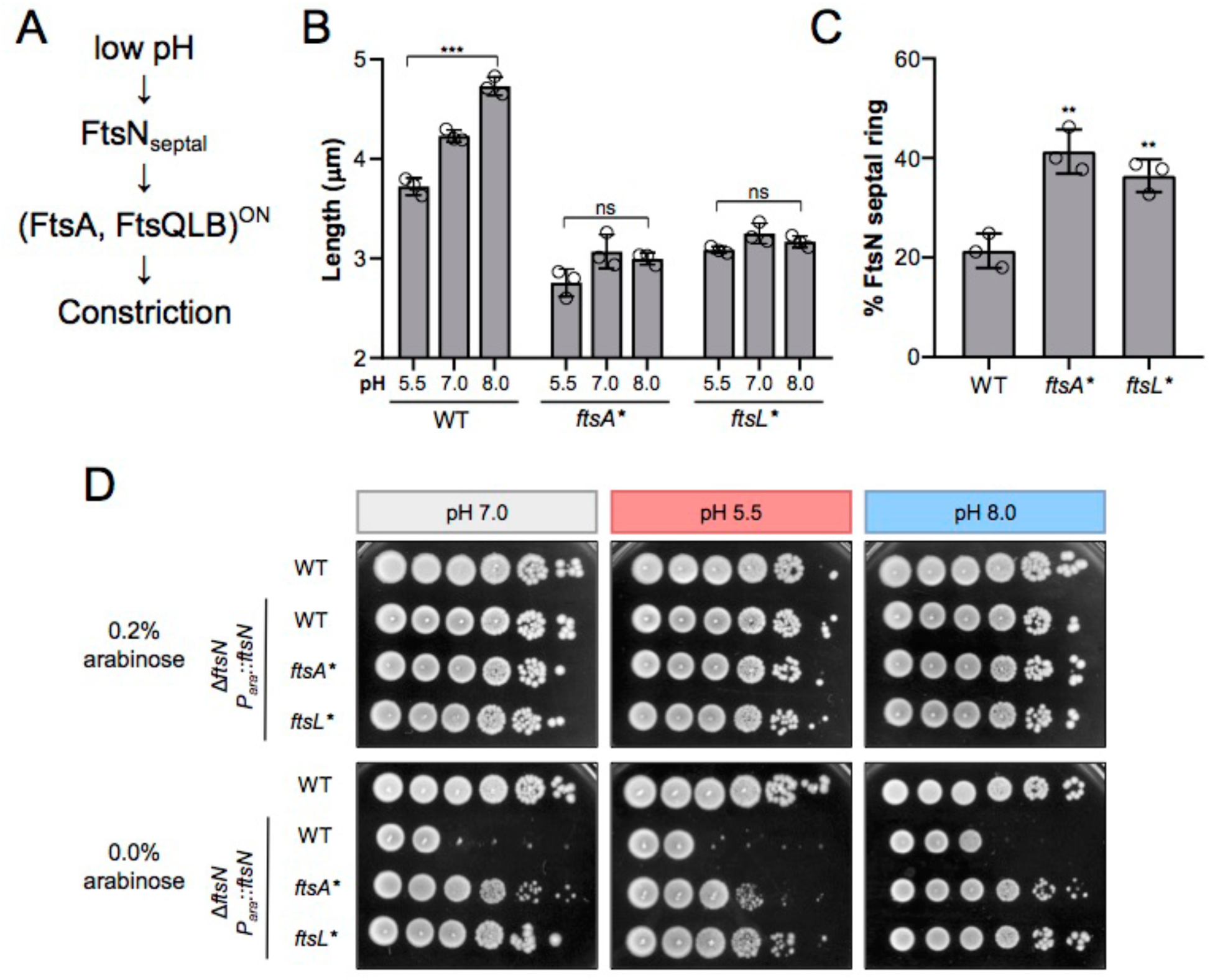
Acidic pH activates division through FtsN. A) Genetic model for pH-dependent activation of cell division. B) Mean cell length of *ftsA\** and *ftsL\** in LB media + 0.2% glucose at pH 5.5, 7.0, and 8.0. MG1655 data from Figure 1C is shown for comparison. Individual points denote mean population measurement for each biological replicate. Error bars represent standard error of the mean. Significance was determined using a two-way ANOVA, corrected for multiple comparisons with Sidak’s test. C) Representative plating efficiency for *ftsN* depletion in WT, *ftsA\**, and *ftsL\** cells at pH 5.5 (middle), 7.0 (left), and 8.0 (right). Image is representative of three biological replicates.

These data are consistent with two models for pH-mediated division activation: 1) growth in acidic media recruits FtsN more efficiently to the septum, causing hyperactivation of FtsA and FtsQLB, or 2) pH influences the activation state of FtsA and FtsQLB independent of FtsN. In the latter model, enhanced FtsN recruitment under acidic conditions may be a consequence of pH-dependent activation of FtsA and/or FtsQLB. To differentiate between these models, we attempted to deplete FtsN in wild type cells in acidic media. If acidic pH activates FtsA and FtsQLB independent of FtsN, we anticipated that less FtsN would be required to sustain growth in acidic media. Consistent with previous work [18,33], *ftsA\** and *ftsL\** mutants tolerated significant depletion of FtsN irrespective of pH environment. However, we were unable to deplete FtsN in wild type cells in any pH condition tested, favoring a model in which acidic pH activates division either through FtsN alone or FtsN together with FtsA and FtsQLB (Figure 5A, D).

### Enrichment of septal FtsN in acidic media requires the constriction control domain but not its glycan binding domain

Although our data indicate FtsN is necessary and sufficient for low pH-mediated division activation, it was not clear which regions of FtsN are required for pH-dependent changes in localization. FtsN’s SPOR domain, CCD, and FtsA interaction interface are all believed to influence in its septal localization [49–51]. To clarify which if any of these interactions are pH sensitive, we compared the septal localization frequency of truncations containing both the FtsA interaction interface and CCD (1-105), only the FtsA interaction interface (1-81), or only the SPOR domain (241-319) fused to GFP at low induction levels. If any of these regions affects FtsN septal recruitment across pH environments, we expect to observe an increase in septal localization in acidic conditions, similar to what we previously observed for the full length GFP-FtsN (Figure 3).

Only the fusion containing both the FtsA interaction domain and the CCD, GFP-FtsN(1-105), exhibited expected pH-dependent changes in septal localization frequency (Figure 6A). Since the FtsA domain is exclusively cytoplasmic, and thus protected from sustained changes in pH [14,52], this finding suggests that CCD-mediated interactions with the cell wall synthesis machinery is pH-responsive. Curiously, the septal localization of the GFP-SPOR domain displayed the opposite pattern from the full length and 1-105 fusion proteins: GFP-SPOR was significantly enriched at the septum in alkaline conditions. In contrast, cells producing GFP-FtsN(1-81) rarely were scored positive for a septal ring under any pH condition, indicating interaction with FtsA in the cytosol likely does not drive pH-dependent changes in recruitment.

**Figure 6.**
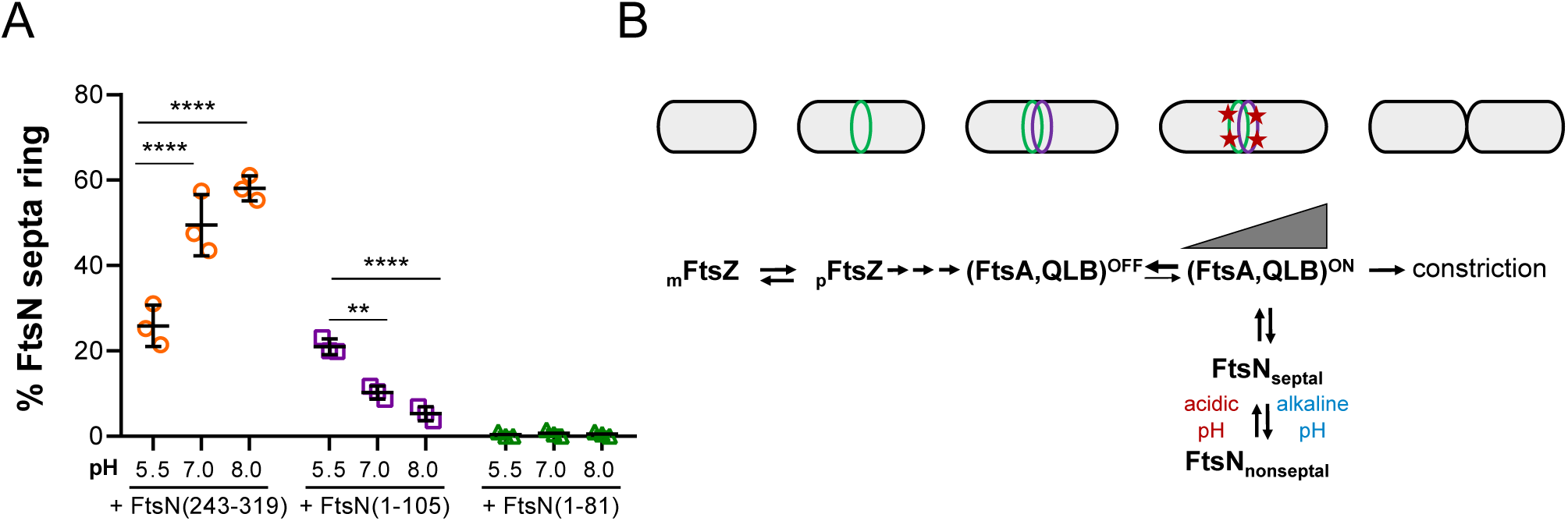
Enrichment of septal FtsN in acidic media requires the CCD but not SPOR domain. A) Mean percentage of cells that score positive for a GFP-FtsN septal ring when producing either the SPOR domain alone (243-319), FtsN(1-105) truncation, or FtsN(1-81) truncation. Cells were grown in LB media and induced with 25 µM IPTG. Individual points depict values of individual biological replicates. Error bars represent standard deviation. Significance was determined using a two-way ANOVA, corrected for multiple comparisons with Tukey’s test. B) Simplified model of divisome activation. Assembly of FtsZ at the septum precipitates recruitment of downstream inactive divisome proteins. Recruitment of FtsN initiates a self-enhancing positive feedback loop of divisome complex activation until a threshold level of activation is achieved, resulting in constriction.

## DISCUSSION

Cytokinesis is a critical integration point for cell size control across all domains of life. In bacteria, cytokinesis consists of three major events: 1) assembly of divisome components to stoichiometric quantities [53], 2) conversion of the machinery into a division competent or “active” state [54], and 3) completion of cross wall synthesis to generate two daughter cells. While previous work focused on the contribution of changes in FtsZ assembly dynamics to cell size [3,6,7], here we identify a central role for the second step, divisome activation, in cell size homeostasis. Specifically, we find that growth in acidic media stimulates cytokinesis in *E. coli* at a smaller volume than that of cells grown in neutral media; conversely, alkaline conditions increase the size at division. Acid-mediated division activation is likely governed by increases in the affinity of the terminal division protein and so-called division “trigger,” FtsN, for other periplasmic components of the cytokinetic ring. An alkaline environment likely has the opposite effect—reducing affinity between divisome components and inhibiting FtsN recruitment. Significantly, pH impacts cell size independent of growth rate, contributing to a growing body of evidence suggesting that these parameters are independent [11,55,56].

### A threshold level of divisome activity dictates cell size

This study illuminates a central role for FtsN in pH-mediated changes in *E. coli* cell size. As the final essential division protein recruited to the septum [31,57], FtsN has long been believed to activate cytokinesis: it interacts with both early and late divisome proteins [18,19,58,59] and its presence at the septum is correlated with visible constriction [32,60,61]. Further supporting a role for FtsN in divisome activation, we find that FtsN septal accumulation is correlated with pH-dependent changes in cell size, and overexpression of FtsN is sufficient to promote cytokinesis at a reduced cell volume independent of media pH (Figure 3, 4). Several additional, less direct pieces of evidence also support FtsN’s involvement in pH-dependent divisome activation. Similar to the phenotypes we observe for cells cultured in acidic media (Figure 2), overexpression of FtsN suppresses the heat sensitivity of cells encoding variants of FtsA, FtsK, FtsQ, and FtsI and bypasses the essentiality of FtsK (SI Appendix, Figure S9) [45,62,63]. We also observed an increase in cell chaining in alkaline pH, particularly during growth in MOPS minimal media (SI Appendix, Figure S1). This is consistent with FtsN’s role in recruiting the septal amidases, which are required for efficient daughter cell separation following division [40,50].

Our data support a refinement of the threshold model for cytokinesis proposed by the Jun lab in which division is coordinated with cell size via growth-rate dependent accumulation of key division proteins to threshold numbers at the future site of septation [3]. Rather than a single mechanism, we favor a “general threshold” model, in which cell division can be coordinated with cell size through disparate if complementary mechanisms. These mechanisms include changes in the specific number of critical divisome proteins at mid-cell as proposed by Si *et al.* and others [3,6,7,17,64,65], alterations in the affinity of divisome proteins for the cytokinetic ring (this work), and changes in the activation state of key regulatory proteins within the ring itself [18,19,66]. Significantly, tuning divisome activity through a variety of mechanisms increases flexibility and allows cells to modulate size in response to a variety of signals. Cytosolic signals (e.g., metabolic state, DNA replication status) may be communicated to the division machinery via regulation of FtsZ or other cytosolic divisome components [6,7], whereas environmental signals that preferentially affect the properties of the periplasm (e.g., pH, osmolarity) could be relayed through differential assembly or activation of the late division machinery rather than through canonical signal transduction cascades.

### A self-enhancing positive feedback loop drives cytokinesis in *E. coli*

Building on previous analyses of interactions between FtsN, the cytoplasmic division protein FtsA, and the periplasmic division proteins FtsQ, FtsL, and FtsB [18,19,51,58,59], our data suggest a mechanism by which environmental pH modulates size. Under physiological conditions, divisome activation requires the conversion of both FtsA and the septal cell wall synthesis machinery into division competent states [54]. FtsN plays a central role in both conversion steps, and its essential activity can be partially or completely bypassed through the expression of hypermorphic alleles of *ftsA*, *ftsL*, *ftsB,* and *ftsW* [18,19,33]. FtsN’s ability to modulate size requires both its FtsA interaction interface and the essential activity of its CCD (Figure 4). The idea that FtsN’s primary role is to activate FtsA and the septal cell wall synthesis machinery is further supported by our finding that the impact of environmental pH on divisome activation and cell size is epistatic to *ftsA\** and *ftsL*.* At the same time, the continued requirement for FtsN at low pH suggests pH-dependent divisome activation is mediated at least in part through FtsN rather than via direct activation of FtsA and FtsQLB (Figure 5).

In contrast to a linear activation pathway in which FtsN localization precedes FtsA and septal cell wall synthesis activation, our findings indicate the presence of a positive feedback loop that builds upon weak interactions between divisome components to generate a mature division complex and initiate cross wall synthesis[19,54] (Figure 6B). We favor a scenario in which divisome proteins are recruited to the future septum in an inactive state where they assemble into discrete complexes. Initial activation of individual complexes, through an as of yet unknown mechanism, stimulates a self-reinforcing feedback loop that enhances FtsN recruitment via interactions with FtsA and FtsQLB until a threshold level of complex activation is achieved and triggers division. Although our data indicate FtsN’s SPOR domain is not required for the initial divisome activation step, this domain is likely to further enhance FtsN recruitment after the onset of constriction when denuded glycans are exposed through the activity of the septal amidases [32,40,50]. The presence of a self-reinforcing feedback loop driving FtsN localization is consistent with previous work indicating synergy between cytoplasmic and periplasmic components of the division machinery with regard to FtsN recruitment. This model is also supported by our finding that 1) FtsN localization enhances the activity of the cytoplasmic and periplasmic components of the division machinery, and 2) activation of FtsA and FtsL reinforce FtsN recruitment to the septum (Figure 5). A self-reinforcing feedback loop requiring iterative rounds of FtsN recruitment provides the cell the opportunity to integrate multiple internal and external signals to ensure that division is tightly coordinated with both cell growth and DNA replication in the face of a hostile and unpredictable environment.

### Environmental sensitivity as a general feature of extracellular processes

While our data favor a specific role for FtsN in driving pH-dependent changes in *E. coli* size, we anticipate that pH has pleiotropic effects on the division machinery. In particular, we speculate that pH affects many, if not all, of the extracellular divisome proteins either by directly impacting activity or through more subtle changes in protein-protein interactions. Consistent with this model, we and others have previously identified a multitude of pH sensitive extracellular cell wall enzymes with diverse enzymatic functions [43,67–69]. *S. aureus* and *S. pneumoniae* still undergo pH-dependent changes in size but lack identifiable homologs of FtsN (SI Appendix, Figure S3) [70], indicating the existence of additional pH-responsive divisome components at least in these organisms. Recent technological advancements, including the use of FRET biosensors to probe interactions between division proteins and new methods to assay activity and interactions between the membrane-associated divisome components, offer promising avenues to dissect the impact of pH specific divisome interactions in future studies [48,71].

More broadly, our results point to the environment as an important mediator of extracytoplasmic processes [72] with pH sensitivity representing just the ‘tip of the iceberg’. Although the underlying molecular mechanisms remain unclear, media osmolarity affects the growth of cells harboring conditional mutants of division genes, dictates essentiality of FtsEX for division, and modulates cell size in *E. coli* [44,73,74]. Extracellular metal availability also alters the activity of several nonessential cell wall enzymes [75–77]. Further exploration promises to illuminate additional environmental determinants modulating the activity of not only enzymes involved in bacterial cell wall synthesis, but also other extracytoplasmic processes.

## MATERIALS AND METHODS

### Bacterial strains, plasmids, and growth conditions

Unless otherwise indicated, all chemicals, media components, and antibiotics were purchased from Sigma Aldrich (St. Louis, MO). Bacterial strains and plasmids used in this study are listed in SI Appendix, Table S1 and S2, respectively. All *E. coli* experiments, with the exception of Figure 5D and SI Appendix Figure S3, were performed in the MG1655 background, referred to as ‘wild type’ in the text. P1 transduction was used to move alleles of interest between strains, and transductants were confirmed with diagnostic PCR and/or sequencing. Mutants were generated using the Q5 Site-Directed Mutagenesis Kit (New England Biolabs). Unless otherwise indicated, *E. coli* strains were grown in lysogeny broth (LB) media (1% tryptone, 1% NaCl, 0.5% yeast extract) with the pH fixed with concentrated NaOH or HCl prior to autoclaving and supplemented with 0.2% glucose. Media pH was confirmed after sterilization. *S. aureus* strains were grown in tryptic soy broth (TSB) with the pH fixed with concentrated NaOH or HCl prior to autoclaving. Where indicated, media was supplemented with 100 mM MES (pH 5.0) or HEPES (pH 7.0 or 8.0) buffers. Cells were cultured in the indicated media at 37 °C shaking at 200 rpm. When selection was necessary, cultures were supplemented with 50 µg/mL kanamycin (Kan), 30 µg/mL chloramphenicol (Cm), 12.5 µg/mL tetracycline (Tet), and/or 25–100 µg/mL ampicillin (Amp).

### Image acquisition

Phase contrast and fluorescence images of fixed cells were acquired from samples on 1% agarose/PBS pads with an Olympus BX51 microscope equipped with a 100X Plan N (N.A. = 1.25) Ph3 objective (Olympus), X-Cite 120 LED light source (Lumen Dynamics), and an OrcaERG CCD camera (Hammamatsu Photonics) or a Nikon TiE inverted microscope equipped with a 100X Plan N (N.A. = 1.25) objective (Nikon), SOLA SE Light Engine (Lumencor), heated control chamber (OKO Labs), and ORCA-Flash sCMOS camera (Hammamatsu Photonics). Filter sets for fluorescence were purchased from Chroma Technology Corporation. Nikon Elements software (Nikon Instruments) was used for image capture.

### Cell size analysis

To achieve balanced growth, cells were cultured from a single colony and grown to exponential phase (OD_600_ ∼ 0.2-0.6). Cultures were then back-diluted into fresh media to an OD_600_ = 0.005 and grown to early exponential phase (OD_600_ between 0.1-0.2) prior to being sampled and fixed for analysis. Cells (500 uL) were fixed by adding 20 µL of 1M NaPO4 (pH 7.4) and 100 µL of fixative (16% paraformaldehyde and 8% glutaraldehyde). Samples were incubated at room temperature for 15 min then on ice for 30 min. Fixed cells were pelleted, washed three times in 1 mL 1X PBS (pH 7.4), then resuspended in GTE buffer (glucose-tris-EDTA) and stored at 4 °C. Images were acquired for analysis within 48 hr of fixation. Cell length, width, and area of *E. coli* cells were determined from phase contrast images using either the ImageJ plugin Coli-Inspector (Figure 1, 5) [47] or the MATLAB software SuperSegger (Figure 4) [78]. For *S. aureus*, cells radii were manually measured and used to calculate cell area. Cell measurements from at least 200 cells from each of at least 3 biological replicates were used to generate single point plots and histograms. Wild type or reference controls were performed during each experiment.

### Time lapse microscopy

Wild type cells in early exponential phase (5 µl) were transferred to a 1% agarose/LB + 0.2% glucose pad at the indicated pH, allowed to dry for 10 minutes, and then imaged on a Nikon TiE inverted microscope heated to 37 °C. Phase contrast images were acquired every 2 minutes for 2 hours. ΔL (change in length) for each cell was calculated in from L_death_ – L_birth_ in SuperSegger [78]. Cells that existed for fewer than 2 frames or more than 20 frames or grew by less than 0.5 uM between birth and death were excluded from the analysis.

### Septal ring frequency analysis

Strains producing GFP fusions were cultured, sampled, and fixed as in the section entitled ‘Cell size analysis.’ Induction conditions for each strain are provided in SI Appendix, Table S4. Phase contrast and fluorescence images were acquired on either a Nikon TiE inverted microscope or Olympus BX51 microscope. The presence of a GFP ‘ring’ for each cell was determined manually: cells were considered positive for a septal ring if they contained a visible band of GFP across the width of the cells or if a single spot of GFP was visible at the midpoint of an invaginating septum. Septal ring frequencies were determined from at least 200 cells from each of at least 3 biological replicates to generate single point plots.

### Heat sensitivity assays

Strains harboring alleles that encode for heat sensitive variants of division proteins were grown in LB (pH 7.0) at 30 °C until mid-log phase (OD_600_ ∼ 0.2-0.6). Cells were pelleted, washed 1x in LB-no salt media, and resuspend in LB-no salt media to an OD_600_ = 1.0. Cells were diluted in LB-no salt media, and serial dilutions 10^-1^ to 10^-6^ were plated under permissive and non-permissive conditions for each mutant with the pH of the plate varying. Plates were incubated for 20 hours. Each experiment was performed at least three times with representative images shown. The permissive condition shown for strains harboring the *ftsZ84*, *ftsA27*, and *ftsQ1* alleles is LB-no salt plates incubated at 30 °C; the non-permissive condition shown for these strains is LB-no salt plates incubated at 37 °C. The permissive conditions shown for strains harboring the *ftsI23* and *ftsK44* alleles are LB-no salt plates incubated at 37 °C; the non-permissive condition shown for these strains is LB-no salt plates incubated at 42 °C.

### Immunoblotting

Strains were grown from a single colony in LB at the indicated pH to mid-log phase (OD_600_ ∼0.2–0.6), back-diluted to 0.005 in 5 mL of media, and grown to an OD_600_ between 0.2–0.3. For experiments measuring native FtsN levels, a MG1655 *malE*::*kan* strain was used to eliminate cross-reactivity with the similarly sized maltose binding protein; this was necessary as antiserum was raised against a FtsN-MBP fusion protein. Samples were pelleted, re-suspended in 2x Laemmli buffer to an OD_600_ ∼20, and boiled for ten minutes. Samples of equivalent volumes were separated on 12% SDS-PAGE gels by standard electrophoresis and transferred to PVDF membranes. Blots were probed with FtsN (1:5000), GFP (1:2000; Abcam), and FtsZ (1:5000) rabbit antiserum and HRP-conjugated secondary antibody (1:5000-1:10000; goat anti-rabbit). Blots were imaged on a LiCor Odyssey imager. Quantitation was determined in FIJI on background subtracted images and normalized to Ponceau staining as a total protein loading control [79].

### FtsN depletion

Strains were grown from a single colony in LB in the presence of inducer (0.2% arabinose) at the indicated pH to mid-log phase (OD_600_ ∼0.2–0.6). Cells were pelleted, washed 3x in LB media (no inducer), and resuspend in LB to an OD_600_ = 1.0. Cells were diluted in LB media, and serial dilutions 10^-2^ to 10^-7^ were plated onto plates with and without inducer (0.2% arabinose) at pH 5.5, 7.0, and 8.0. Plates were incubated for 20 hours. Each experiment was performed at least three times with representative images shown.

### Statistical analysis

A minimum of three biological replicates were performed for each experimental condition unless otherwise indicated. Data are expressed as means ± standard deviation (SD) or standard error of the mean (SE). Statistical tests employed are indicated in the text and corresponding figure legend. Analysis was performed in GraphPad Prism. No statistical methods were used to predetermine sample size. Asterisks indicate significance as follows: *, p<0.05; **, p<0.01; ***, p<0.001; ****, p<0.0001.

## ACKNOWLEDGEMENTS

We thank David Weiss, Thomas Bernhardt, Piet de Boer, Bill Margolin, and Joe Lutkenhaus for kind gifts of strains, plasmids, and antibodies necessary to carry out this work. We thank David Weiss and members of the Levin lab for helpful discussions. This work was supported by NIH GM127331 (P.A.L.), an Arnold O. Beckman postdoctoral fellowship (C.S.W.), NSF graduate research fellowship DGE-1745038 (E.A.M.), and a Center for Science and Engineering of Living Systems graduate scholar fellowship (E.A.M.).

