## Supplemental Information for "Environmental pH impacts division assembly and cell size in *Escherichia coli*"

**This PDF file includes:**

SI Methods
Figures S1 to S12
Tables S1 to S5
SI References

### SI METHODS

#### Mid-cell intensity quantification

Population demographs from at least 1000 cells from at least three biological replicates were generated in Coli Inspector [1]. Mid-cell intensity as a function of cell length was determined by manually drawing a line (line width = 6) through the mid-cell and plotting the line profile in FIJI [2]. Background intensity, the average of two non-mid cell sites, was subtracted from each data point. The data was smoothed and analyzed in GraphPad prism.

### SUPPLEMENTAL FIGURES

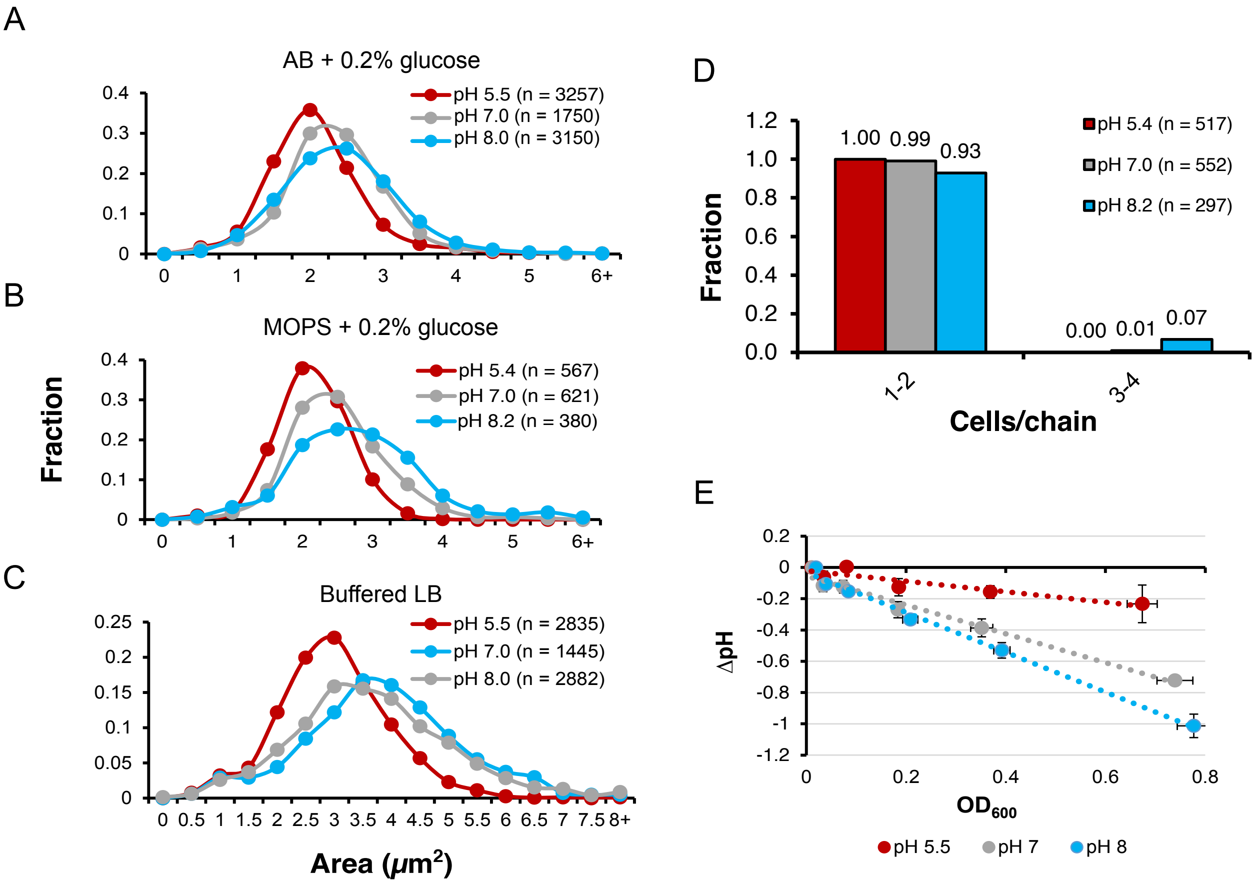


**Figure S1.** pH-dependent changes in cell size are independent of growth medium and buffering capacity.
A-C) Histograms of MG1655 cell area distribution for cells grown in AB minimal media + 0.2% glucose (A), MOPS minimal media + 0.2% glucose (B), or LB media supplemented with 100 mM MES (pH 5.5), HEPES (pH 7.0 or pH 8.0) (C).
D) Fraction of cells present in chains as a function of media pH during growth in MOPS minimal media + 0.2% glucose.
E) Change in pH as a function of optical density in unbuffered LB media. Cells were inoculated at an $OD_{600}$ = 0.005.

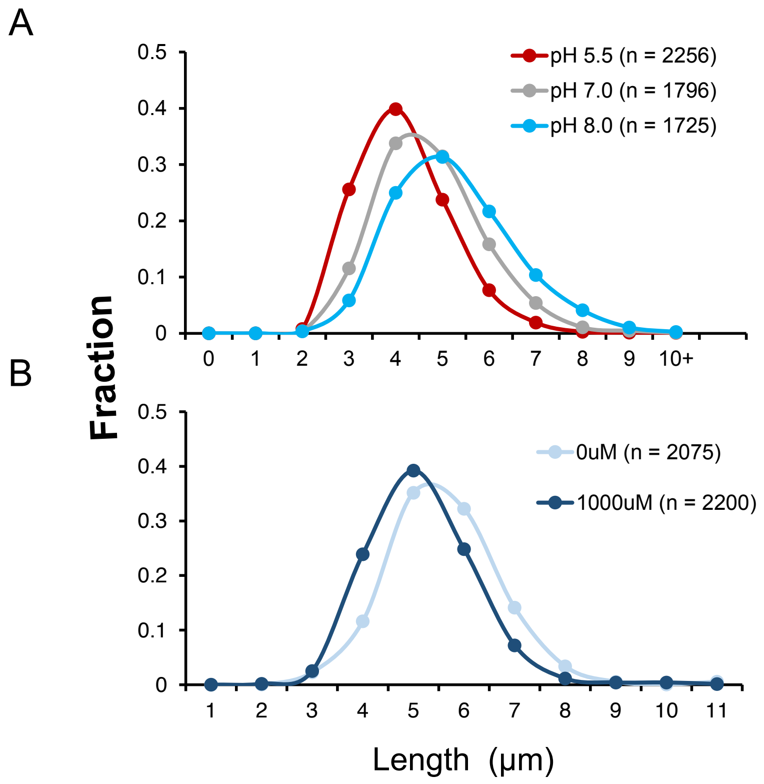


**Figure S2.** Distribution of MG1655 cell lengths as a function of pH (A) or FtsN overexpression from pCH201 plasmid (B).

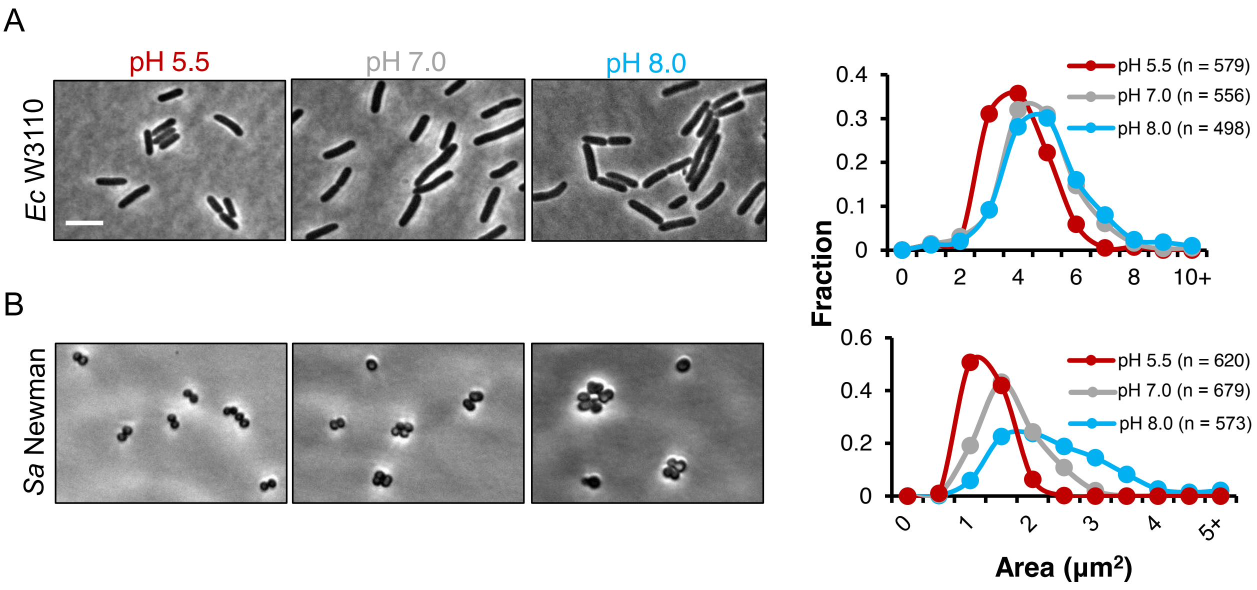


**Figure S3.** Evolutionarily distant bacteria undergo pH-dependent changes in cell size.
A-B) Representative micrographs and cell area distributions for *E. coli* strain W3110 grown in LB + 0.2% glucose (A) and *S. aureus* strain Newman grown in TSB (B) as a function of environmental pH. Scale bar denotes 5 µm.

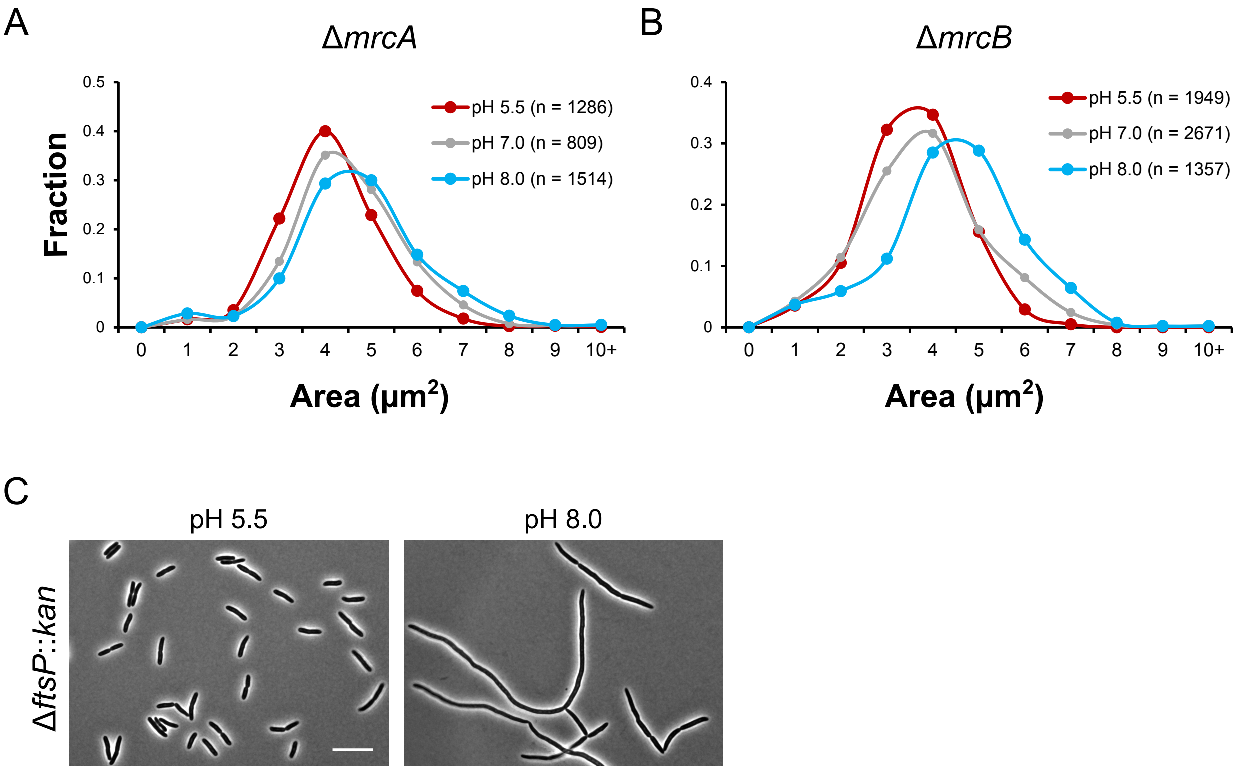


**Figure S4.** Accessory divisome factors do not participate in pH-dependent changes in cell size.
A-B) Cell area distributions for MG1655 strains defective for PBP1a (A) and PBP1b (B) production as a function of pH during growth in LB + 0.2% glucose.
C) Representative micrographs of MG1655 strain defective for FtsP during growth in LB + 0.2% glucose at pH 5.5 (left) and pH 8.0 (right).

A

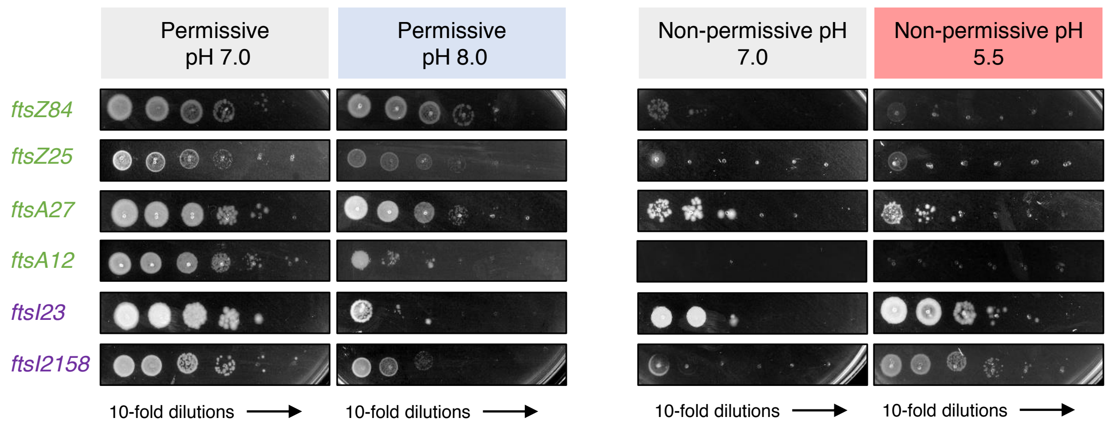


B

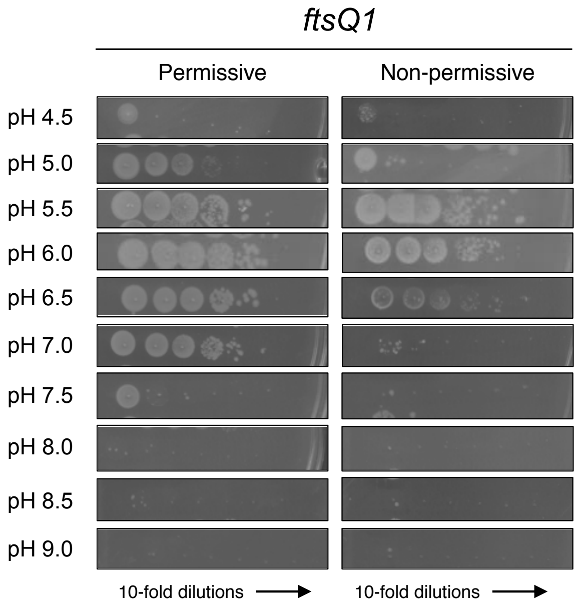


C

| **pH** | **Permissive, pH 8.0** | | | **Non-permissive, pH 5.5** | | |
|---|---|---|---|---|---|---|
| | *ftsQ1* | *ftsK44* | *ftsI23* | *ftsQ1* | *ftsK44* | *ftsI23* |
| 4.5 | ** | - | - | - | - | - |
| 5.0 | - | - | - | - | ++ | ++ |
| 5.5 | - | - | - | ++ | ++ | ++ |
| 6.0 | - | - | - | ++ | + | + |
| 6.5 | - | - | - | ++ | - | - |
| 7.0 | - | - | - | - | - | - |
| 7.5 | ** | * | * | - | - | - |
| 8.0 | ** | ** | * | - | - | - |
| 8.5 | ** | ** | ** | - | - | - |
| 9.0 | ** | ** | ** | - | - | - |

**Figure S5.** Mutants producing temperature sensitive late division proteins are suppressed in acidic conditions and enhanced in alkaline conditions.
A) Representative plating efficiency for cells producing unique temperature sensitive variants of FtsZ, FtsA, and FtsI during growth at permissive (left) or non-permissive (right) condtions.
B) Representative plating efficiency for cells harboring the *ftsQ1* allele upon exposure to a wide pH range under permissive (left) and non-permissive (right) conditions.
C) Table summarizing suppression and enhancement data for strains harboring temperature sensitive variants in late division proteins across a range of pH conditions. ++, +, and – denote complete, partial, or no suppression at the indicated pH. **, *, and – denote complete, partial, or no enhancement at the indicated pH.

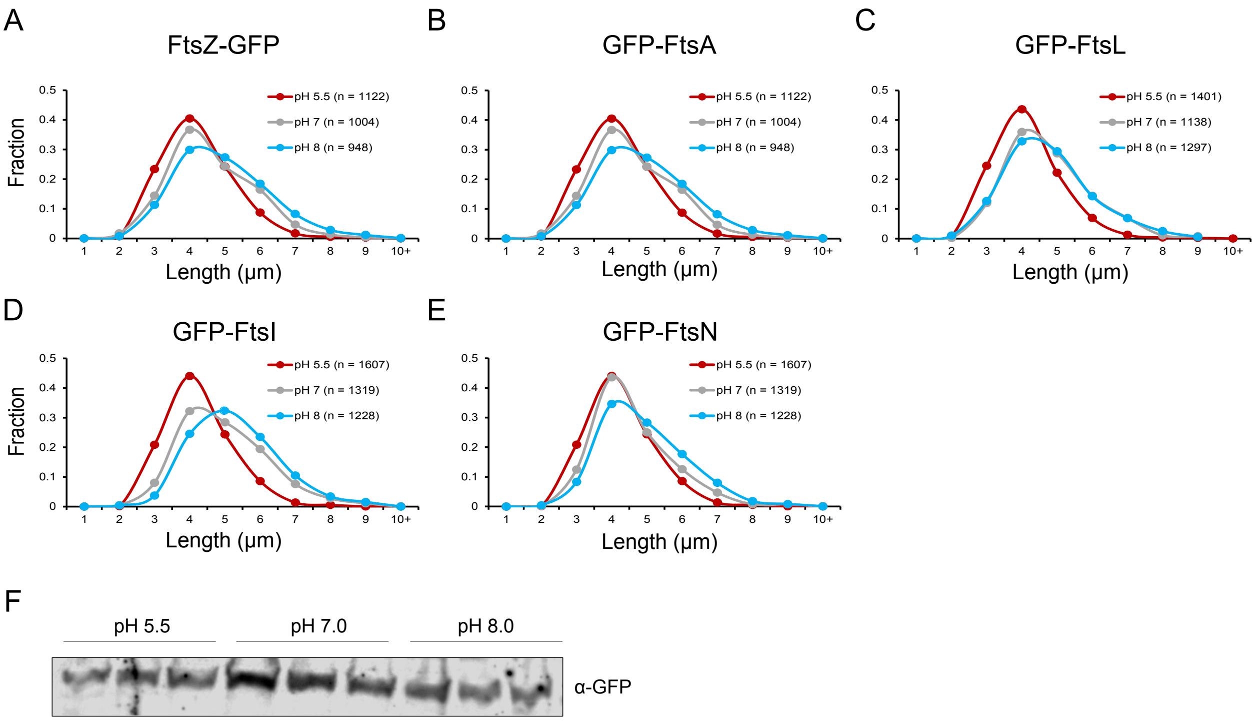


**Figure S6.** Production of GFP-tagged division proteins does not eliminate pH-dependent changes in cell length.
A-E) Cell length distributions of cells overexpressing GFP-tagged division proteins at pH 5.5, 7.0, and 8.0.
F) Western blot for GFP-FtsN levels in EAM621 as a function of pH. Three replicates for each pH condition are shown.

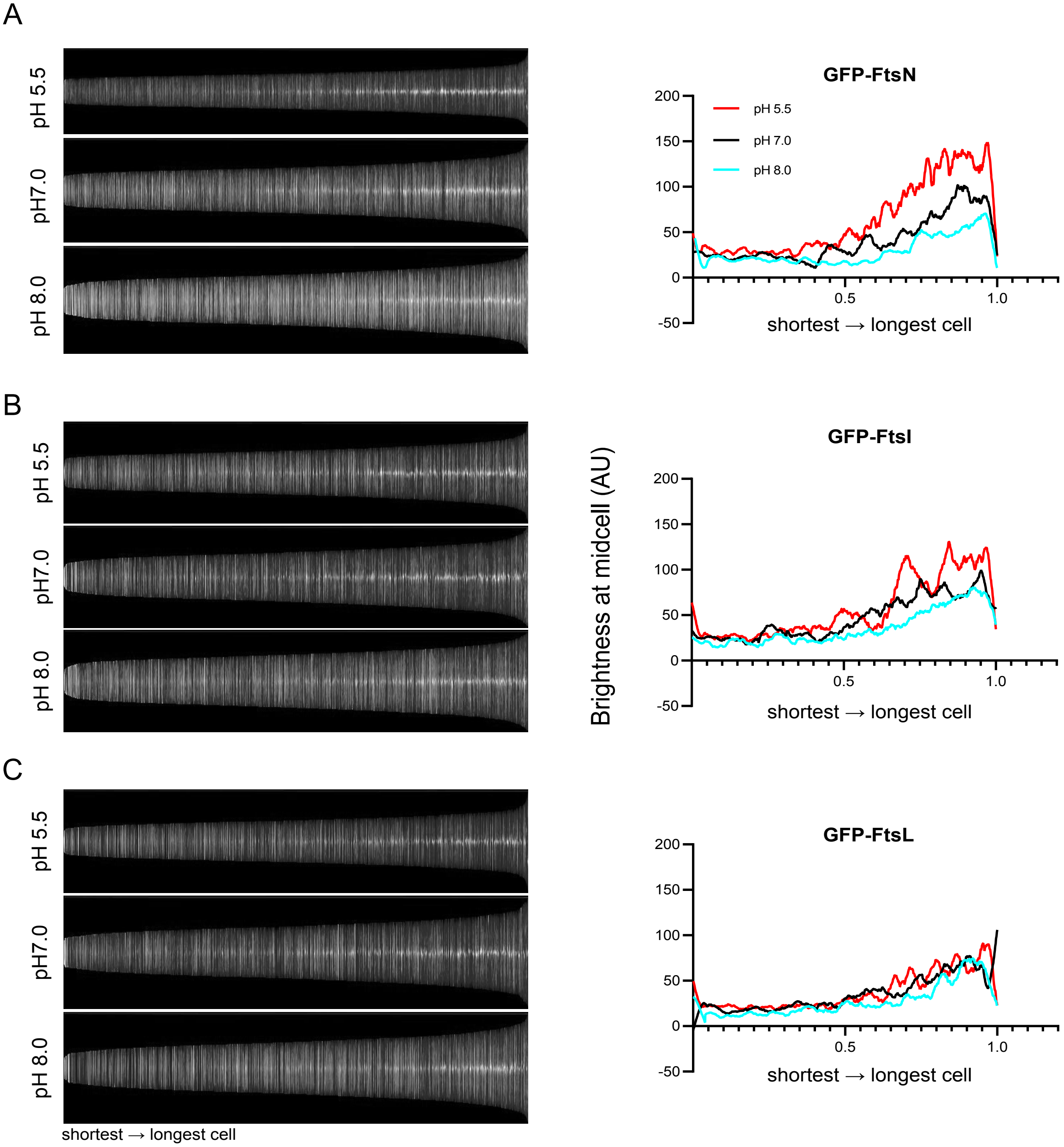


**Figure S7.** Intensity of septal localization of GFP-tagged late division proteins across pH conditions. A-C) Demographs (left) and mid-cell intensity quantifications (right) for cells producing GFP-FtsN (A), GFP-FtsI, and GFP-FtsL (C). At least 1000 were analyzed for each strain and pH condition.

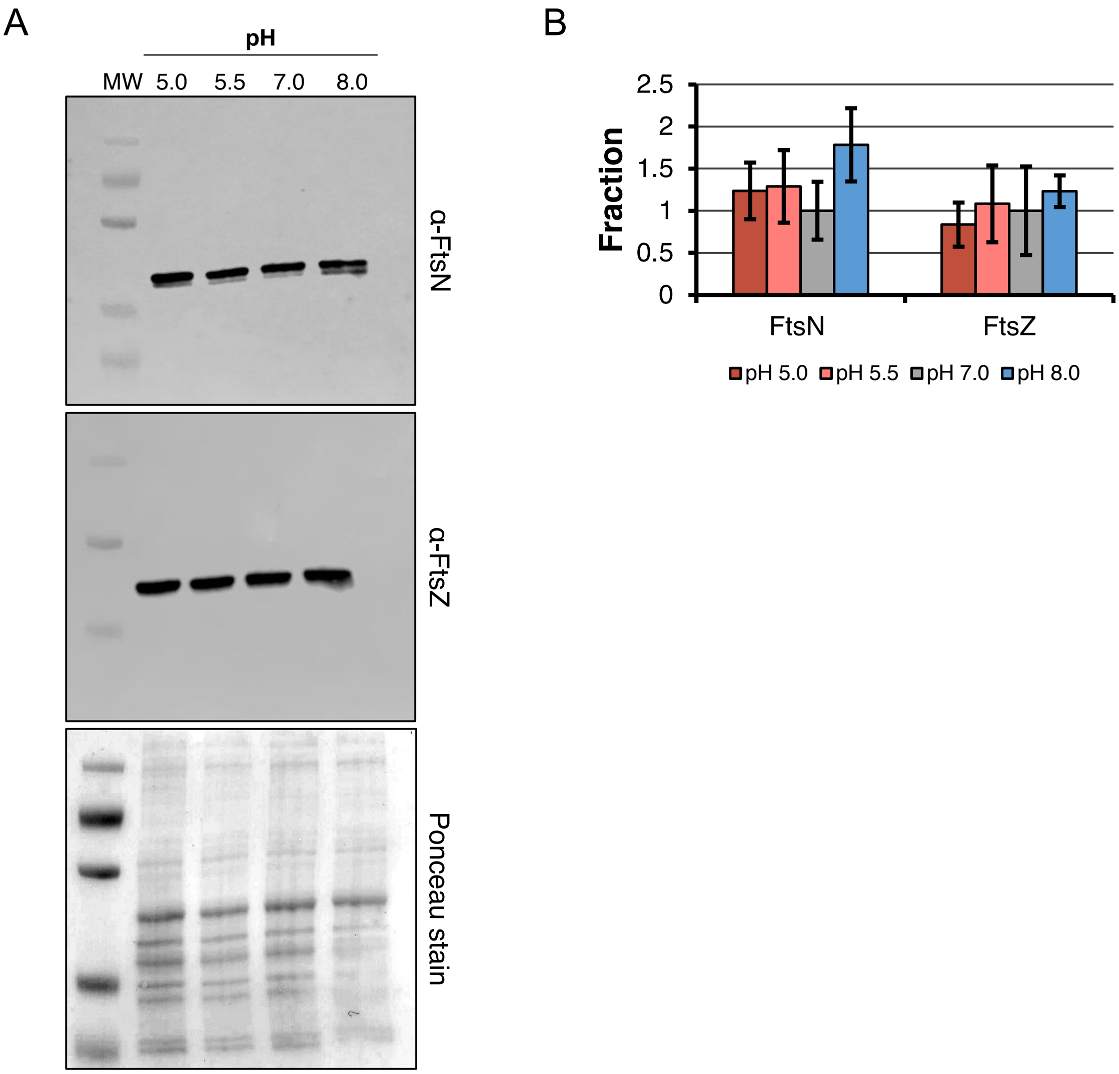


**Figure S8.** Production of FtsN does not vary across pH conditions.
A) Uncropped membrane shown in Figure 3 probed with anti-FtsN sera (top) and anti-FtsZ sera (middle) or strained with Ponceau reagent for total protein levels (bottom).
B) Quantification of relative FtsN and FtsZ levels as a function of pH. Bars depict mean relative levels of each protein ± SD relative to pH 7.0 from three independent cultures and normalized for total protein load as determined by Ponceau stain.

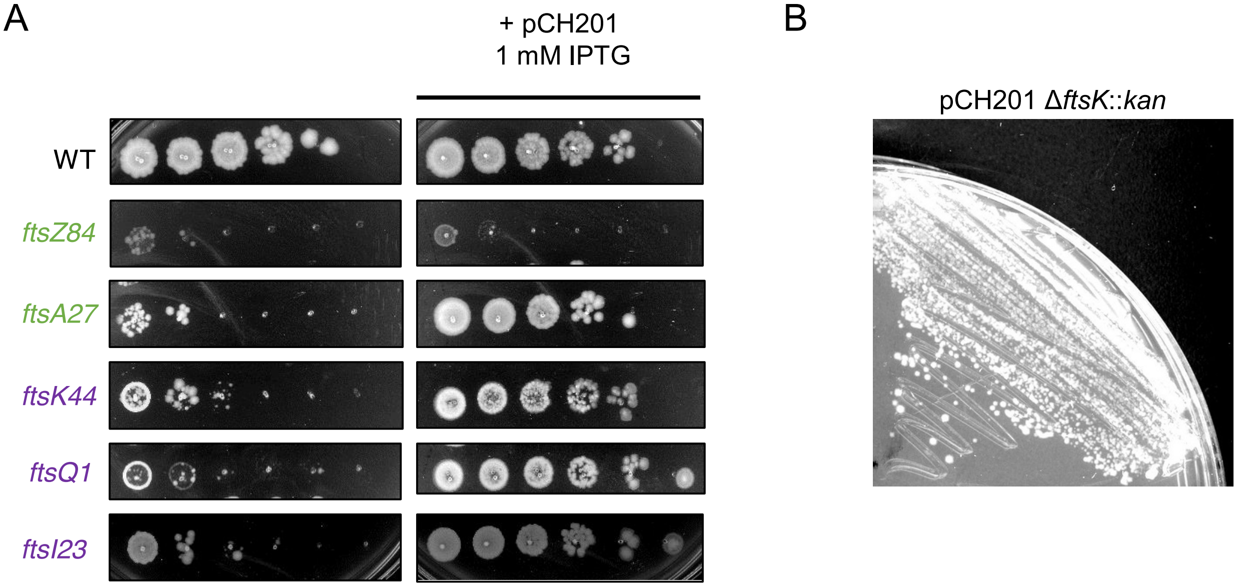


**Figure S9.** FtsN overexpression suppresses a subset of temperature sensitive division alleles and bypasses the essential function of FtsK.
A) Representative plating efficiency for cells producing temperature sensitive variants of division proteins under non-permissive growth conditions in the presence (right) or absence (left) of FtsN overexpression.
B) MG1655 can grow in the absence of FtsK upon FtsN overexpression (1 mM IPTG).

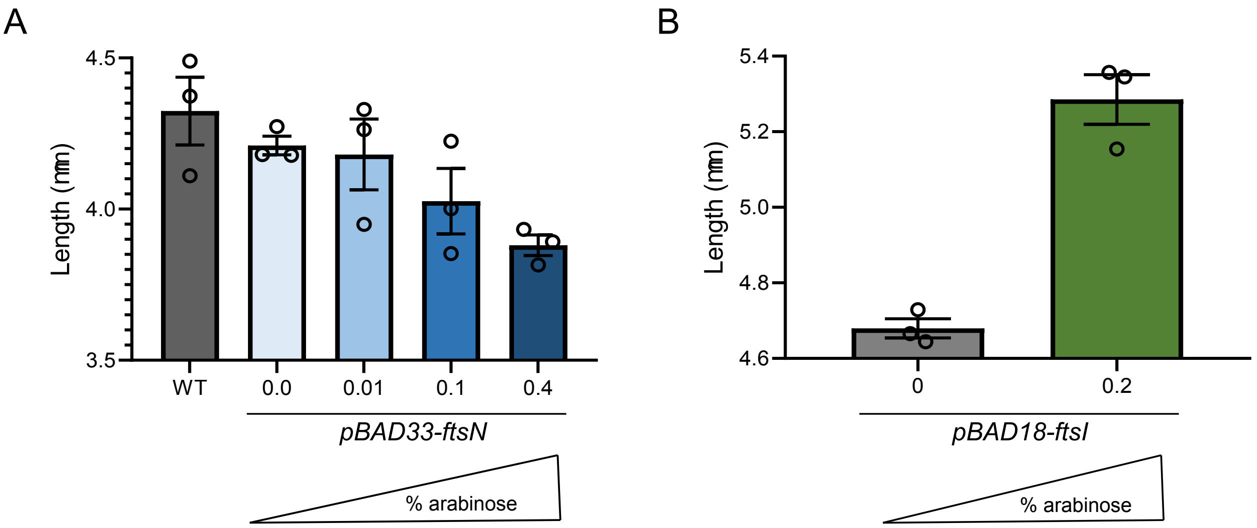


**Figure S10.** Impact of late division protein overexpression on cell length.
A-B) Cell length of MG1655 producing excess FtsN (A) or FtsI (B) during growth in LB media. Bars represent mean cell length ± SEM from three independent biological replicates (n > 200 cells per replicate).

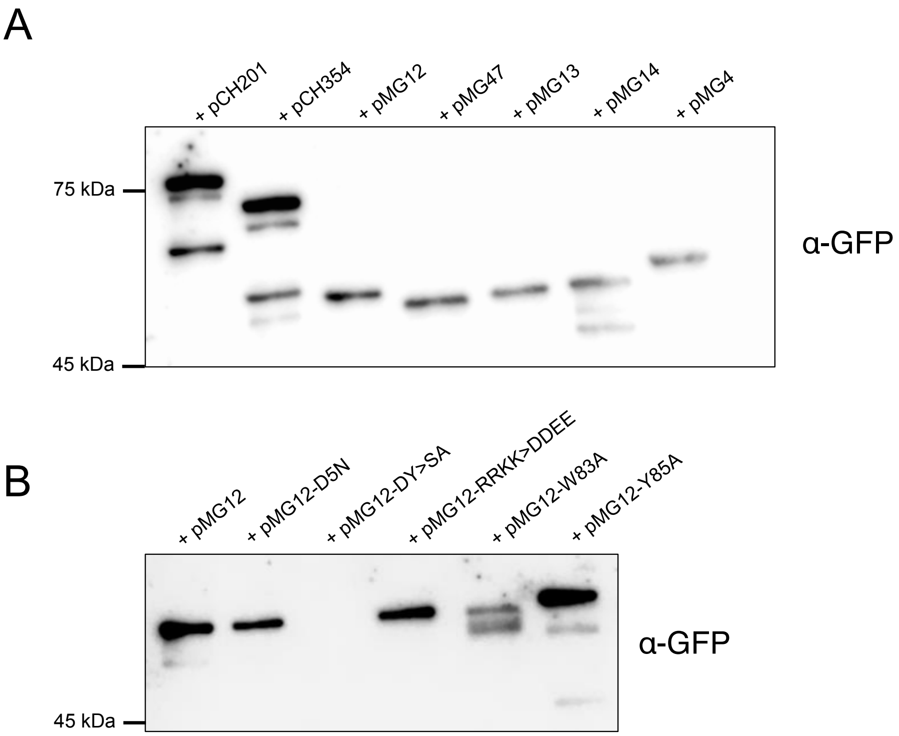


**Figure S11.** Production of GFP-FtsN variants.
A-B) Representative Western blots for GFP-FtsN truncations (A) or point mutants (B) expressed in MG1655 (+1 mM IPTG) and probed with anti-GFP.

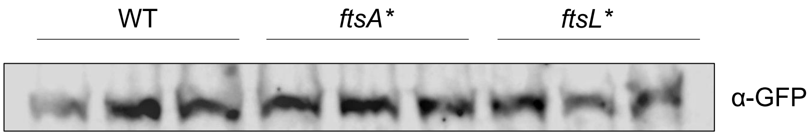


**Figure S12.** Production of GFP-FtsN does not vary in division hypermorph mutants.
Western blot depicting GFP-FtsN levels from MG1655, *ftsA**, and *ftsL** grown in LB + 0.2% glucose media (pH 7.0). Three biological replicates are shown for each strain.

### SUPPLEMENTAL TABLES

**Table S1.** Bacterial strains used in this study.

| **Designation** | **Genotype** | **Source**[a] |
|---|---|---|
| MG1655 (*E. coli*) | *rph1 ilvG rfb-50 λ- F-* | [3] |
| W3110 (*E. coli*) | *rph1 IN(rrnD-rrnE)1 λ- F-* | [4] |
| MC4100 (*E. coli*) | *araD129 ΔlacU169 relA1 rpsL150 thi mot flb5301 deoC7 rbsR F-* | [5] |
| Newman (*S. aureus)* | | [6] |
| TB28 | MG1655 *lacIZYA*::*frt* | [7] |
| EAM696 | MG1655 *mrcB*::*frt* | [8] |
| EAM899 | MG1655 *mrcA*::*frt* | [8] |
| EAM1081 | MG1655 *ftsP*::*kan* | P1(JW2985[b]) x MG1655 |
| CW142 | MG1655 *malE*::*kan* | P1(JW3994[b]) x MG1655 |
| BH330 | MG1655 $P_{lac}$-*gfp-ftsZ* | [9] |
| EC479 | MC4100 $P_{210}$-*gfp-ftsA leu*::Tn*10* | [10] |
| EAM410 | MG1655 $P_{210}$-*gfp-ftsA leu*::Tn*10* | P1(EC479) x MG1655 |
| PAL3700 | TB28 $P_{lac}$-*gfp-ftsL* | [11] |
| EC454 | MC4100 $P_{207}$-*gfp-ftsI leu*::Tn*10* | [10] |
| EAM412 | MG1655 $P_{207}$-*gfp-ftsI leu*::Tn*10* | P1(EC454) x MG1655 |
| EAM621 | MG1655 $P_{204}$-*gfp-ftsN* | [12] |
| PAL2452 | MG1655 *leu82*::Tn*10 ftsZ84* | [13] |
| WM4649 | MG1655 *lacU169 leu82*::Tn*10 ftsI23* | [14] |
| EC433 | MG1655 *leu82*::Tn*10 ftsQ1* | [15] |
| MM61 | MG1655 *leu82*::Tn*10 ftsA12* | [15] |
| WM2101 | MG1655 *lacU169 ycaD::*Tn*10 ftsK44* | [16] |
| WM4107 | MG1655 *lacU16 leu-260*::Tn*10 ftsA27* | [17] |
| PAM161 | *ftsZ25* | [18] |
| AX655 | *ftsI2158* | [19] |

| | | |
|---|---|---|
| BH142 | MG1655 *leu82*::Tn*10 ftsA** | [20] |
| MT13 | TB28 *leu82*::Tn*10 ftsL** | [11] |
| EAM747 | MG1655 *leu82*::Tn*10 ftsA** $P_{204}$*-gfp-ftsN* | P1(EAM621) x BH142 |
| EAM749 | TB28 *leu82*::Tn*10 ftsL** $P_{204}$*-gfp-ftsN* | P1(EAM621) x MT13 |
| MT75 | TB28 *ftsK*::*kan* | [11] |
| EAM1311 | MG1655 *ftsK*::*kan* | P1(MT75) x MG1655 |
| HSC074 | MC4100 *ftsN*::*kan* | [21] |
| EAM719 | MC4100 *ftsN*::*kan leu82*::Tn*10 ftsA** | P1(BH142) x HSC074 |
| EAM723 | MC4100 *ftsN*::*kan leu82*::Tn*10 ftsL** | P1(MT13) x HSC074 |

[a] Strains constructed by P1 transduction are described using the shorthand: P1(donor) x recipient.
[b] Strains sourced from the Coli Genetic Stock Center [22]

**Table S2.** Plasmids used in this study

| Designation | Genotype | Source |
|---|---|---|
| pCH201 | *bla* $lacI^q$ $P_{lac}$::*gfp-FtsN(1-319)* | [23] |
| pCH354 | *bla* $lacI^q$ $P_{lac}$::*gfp-FtsN(1-243)-le* | [23] |
| pMG12 | *bla* $lacI^q$ $P_{lac}$::*gfp-FtsN(1-105)-le* | [23] |
| pMG47 | *bla* $lacI^q$ $P_{lac}$::*gfp-FtsN(1-90)* | [23] |
| pMG13 | *bla* $lacI^q$ $P_{lac}$::*gfp-FtsN(1-81)-le* | [23] |
| pMG14 | *bla* $lacI^q$ $P_{lac}$::$^{ss}$*torA-gfp-FtsN(71-105)-le* | [23] |
| pMG4 | *bla* $lacI^q$ $P_{lac}$::$^{ss}$*torA-gfp-FtsN(341-319)-le* | [23] |
| pMG12-D5N | *bla* $lacI^q$ $P_{lac}$::*gfp-FtsN(1-105)-le(D5N)* | This work |
| pMG12-RRKK>DDEE | *bla* $lacI^q$ $P_{lac}$::*gfp-FtsN(1-105)-le(RRKK>DDEE)* | This work |
| pMG12-W83A | *bla* $lacI^q$ $P_{lac}$::*gfp-FtsN(1-105)-le(W83A)* | This work |
| pMG12-Y85A | *bla* $lacI^q$ $P_{lac}$::*gfp-FtsN(1-105)-le(Y85A)* | This work |
| pBAD33-ftsN | pBAD33-ftsN | [24] |
| pLMG173 | pBAD18-ftsI | [25] |

**Table S3.** Impact of pH on cell dimensions of MG1655 in LB media[a]

| pH | Area ($\mu m^2$)[a,c] | Length ($\mu m$)[a,c] | Width ($\mu m$)[a,c] | MDT (min)[b,c] | *n* cells |
|---|---|---|---|---|---|
| 4.5 | 2.75 ± 0.07 (****) | 3.42 ± 0.12 (****) | 0.85 ± 0.01 (*) | 32 ± 2 (****) | 2600 |
| 5.0 | 2.93 ± 0.07 (***) | 3.45 ± 0.04 (****) | 0.89 ± 0.01 (ns) | 25 ± 1 (***) | 2299 |
| 5.5 | 3.25 ± 0.05 (*) | 3.72 ± 0.05 (***) | 0.92 ± 0.01 (ns) | 22 ± 1 (ns) | 2256 |
| 6.0 | 3.33 ± 0.02 (ns) | 3.92 ± 0.02 (*) | 0.89 ± 0.01 (ns) | 22 ± 1 (ns) | 1588 |
| 6.5 | 3.58 ± 0.14 (ns) | 4.07 ± 0.09 (ns) | 0.91 ± 0.02 (ns) | 21 ± 1 (ns) | 1526 |
| 7.0 | 3.75 ± 0.06 | 4.23 ± 0.04 | 0.92 ± 0.01 | 21 ± 1 | 1796 |
| 7.5 | 3.78 ± 0.15 (ns) | 4.34 ± 0.08 (ns) | 0.90 ± 0.03 (ns) | 21 ± 1 (ns) | 1233 |
| 8.0 | 4.25 ± 0.12 (*) | 4.73 ± 0.05 (***) | 0.93 ± 0.02 (ns) | 21 ± 1 (ns) | 1725 |
| 8.5 | 4.44 ± 0.19 (**) | 4.93 ± 0.08 (****) | 0.93 ± 0.03 (ns) | 22 ± 1 (ns) | 1298 |

[a] ± SEM
[b] ± SD
[c] Statistical significance compared to pH 7.0 indicated in parenthesis as determined by a one-way ANOVA corrected for multiple comparisons with Dunnett's test.

**Table S4.** Septal ring frequencies across pH conditions in LB media.

| Strain/plasmid | [IPTG] (μm) | pH 5.5 ring frequency (%)[a] | pH 7.0 ring frequency (%)[a] | pH 8.0 ring frequency (%)[a] |
|---|---|---|---|---|
| BH300 | 1000 | 87 ± 4 | 86 ± 2 | 90 ± 5 |
| EAM410 | 100 | 81 ± 2 | 85 ± 3 | 87 ± 3 |
| PAL3700 | 100 | 29 ± 6 | 29 ± 2 | 27 ± 6 |
| EAM412 | 2.5 | 27 ± 2 | 26 ± 7 | 23 ± 3 |
| EAM621 | 5 | 31 ± 2 | 23 ± 2 | 14 ± 3 |
| MG1655/pMG4 | 25 | 26 ± 5 | 50 ± 7 | 58 ± 3 |
| MG1655/pMG12 | 25 | 21 ± 2 | 10 ± 2 | 5 ± 2 |
| MG1655/pMG13 | 25 | 0 ± 1 | 1 ± 1 | 1 ± 1 |

[a] ± SD

**Table S5.** Impact of FtsN overexpression on cell size in LB media[a]

| Plasmid | [IPTG] (uM) | Area ($\mu m^2$)[b] | Length (µm)[b] | Width (µm)[b] | MDT (min)[c] | Ring frequency (%)[c,d] | *n* cells |
|---|---|---|---|---|---|---|---|
| N/A | N/A | 3.81 ± 0.11 | 4.23 ± 0.07 | 0.89 ± 0.02 | 22 ± 2 | N/A | 7226 |
| pCH201 | 0 | 3.89 ± 0.10 | 4.15 ± 0.08 | 0.93 ± 0.01 | 23 ± 2 | 0.7 ± 0.3 | 2075 |
| pCH201 | 10 | 3.86 ± 0.10 | 4.08 ± 0.04 | 0.94 ± 0.02 | 22 ± 2 | 6.3 ± 2.0 | 1851 |
| pCH201 | 100 | 3.63 ± 0.10 | 3.80 ± 0.05 | 0.95 ± 0.01 | 23 ± 3 | 34.6 ± 4.5 | 1859 |
| pCH201 | 1000 | 3.42 ± 0.11 | 3.69 ± 0.05 | 0.91 ± 0.02 | 23 ± 2 | 36.6 ± 7.7 | 8036 |
| pCH354 | 1000 | 3.63 ± 0.11 | 3.91 ± 0.15 | 0.92 ± 0.02 | 20 ± 1 | N.D[e] | 2285 |
| pMG12 | 1000 | 3.51 ± 0.11 | 3.88 ± 0.07 | 0.89 ± 0.03 | 21 ± 2 | 10.2 ± 1.6 | 6170 |
| pMG47 | 1000 | 3.82 ± 0.15 | 4.24 ± 0.06 | 0.89 ± 0.04 | 20 ± 1 | N.D | 1864 |
| pMG13 | 1000 | 3.98 ± 0.14 | 4.30 ± 0.11 | 0.91 ± 0.03 | 21 ± 1 | 0.7 ± 0.8 | 1835 |
| pMG14 | 1000 | 4.01 ± 0.26 | 4.44 ± 0.13 | 0.89 ± 0.03 | 21 ± 1 | N.D | 2057 |
| pMG4 | 1000 | 4.18 ± 0.38 | 4.71 ± 0.10 | 0.87 ± 0.07 | 23 ± 3 | 49.5 ± 7.2 | 2778 |
| pMG12-D5N | 1000 | 3.71 ± 0.26 | 4.12 ± 0.18 | 0.89 ± 0.05 | 21 ± 2 | N.D | 3575 |
| pMG12-RRKK>DDEE | 1000 | 4.22 ± 0.02 | 4.40 ± 0.06 | 0.94 ± 0.01 | 21 ± 1 | N.D | 1066 |
| pMG12-W83A | 1000 | 3.79 ± 0.13 | 4.27 ± 0.18 | 0.88 ± 0.06 | 22 ± 1 | N.D | 3083 |
| pMG12-Y85A | 1000 | 3.63 ± 0.29 | 4.16 ± 0.17 | 0.86 ± 0.05 | 22 ± 1 | N.D | 3470 |

[a] All plasmids were transformed into the parental strain MG1655
[b] ± SEM

[c] ± SD
[d] pH 7.0
[e] N.D, not determined
